# Comparing diversification rates in lakes, rivers, and the sea

**DOI:** 10.1101/2020.05.24.113761

**Authors:** Elizabeth Christina Miller

## Abstract

The diversity of species inhabiting freshwater relative to marine habitats is striking, given that freshwater habitats encompass <1% of Earth’s water. The most commonly proposed explanation for this pattern is that freshwater habitats are more fragmented than marine habitats, allowing more opportunities for allopatric speciation and thus increased diversification rates in freshwater. However, speciation may be generally faster in sympatry than in allopatry, as illustrated by lacustrine radiations such as African cichlids. Differences between rivers and lakes may be important to consider when comparing diversification broadly among freshwater and marine lineages. Here I compared diversification rates of teleost fishes in marine, riverine and lacustrine habitats. I found that lakes had faster speciation and net diversification rates than other aquatic habitats. However, most freshwater diversity arose in rivers. Surprisingly, riverine and marine habitats had similar rates of net diversification. Biogeographic models suggest that lacustrine habitats are evolutionary unstable, explaining the dearth of lacustrine species in spite of their rapid diversification. Collectively, these results suggest that diversification rate differences are unlikely to explain the freshwater paradox. Instead, this pattern may be attributable to the comparable amount of time spent in riverine and marine habitats over the 200-million-year history of teleosts.

## INTRODUCTION

The great biodiversity of freshwater habitats is striking. Habitable freshwater makes up 0.009% of the Earth’s water volume but contain at least 40% of ray-finned fish species (Cohen 1970; Horn 1972), a pattern called the “freshwater paradox” (Tedesco et al. 2017a). The diversity of freshwater relative to marine fishes is thought to be due to the greater fragmentation and isolation of freshwater habitats, creating more opportunities for geographic population subdivision than in the ocean (Cohen 1970; Horn 1972; May 1994; Grosberg et al. 2012). For this reason, speciation and net diversification rates are expected to be higher in freshwater than in marine lineages (e.g., Tedesco et al. 2017a).

Two innovations have allowed researchers to rigorously test longstanding ideas about the origin of biodiversity: the expansion in size and taxonomic scope of molecular phylogenies (Rabosky et al. 2013; Rabosky et al. 2018), and the development of increasingly sophisticated approaches to estimate diversification rates (Maddison et al. 2007; Rabosky 2014). The hypothesis that freshwater fishes have faster diversification rates than marine fishes has been tested in at least 10 studies over the past decade, with mixed results (Table 1). Most studies focusing on a single clade of fishes spanning both marine and freshwater biomes have found little difference in net diversification rates (speciation minus extinction rates; see references in Table 1). This could perhaps be due to low power to detect diversification rate differences within small clades (Davis et al. 2013), or because speciation and extinction rates are both higher in freshwater leading to small differences in net diversification (Bloom et al. 2013). Note that higher turnover in freshwater compared to marine habitats cannot explain the freshwater paradox, because species richness will not change when speciation and extinction rates are about equal (see also Meseguer and Condamine 2020). Studies including all ray-finned fishes have found no difference in diversification rates (Carrete Vega and Wiens 2012), faster rates in marine habitats (Betancur-R et al. 2015), faster rates in freshwater (Tedesco et al. 2017a), or faster rates in only some freshwater clades (Rabosky 2020). The only study to my knowledge that exclusively used the fossil record to test this hypothesis found that the accumulation of family diversity has slowed over time in marine but not freshwater fishes (Guinot and Cavin 2015). However, familial diversity is lower overall in freshwater, so it is unknown if freshwater diversity will also plateau if it ever reaches a similar magnitude as the ocean (Figure 1 of Guinot and Cavin 2015). Studies of organisms other than ray-finned fishes are also collectively unclear as to whether marine, freshwater or terrestrial habitats have the fastest diversification rates (Table 1). In sum, whether freshwater or marine fishes have faster diversification rates could be influenced by choice of methods, focal clade, or the breadth of clades included in the study. This sensitivity could exist because the difference in rates between biomes is actually narrower than expected (and therefore hard to detect), or because the habitat with the faster rate differs among clades.

**Figure 1.**
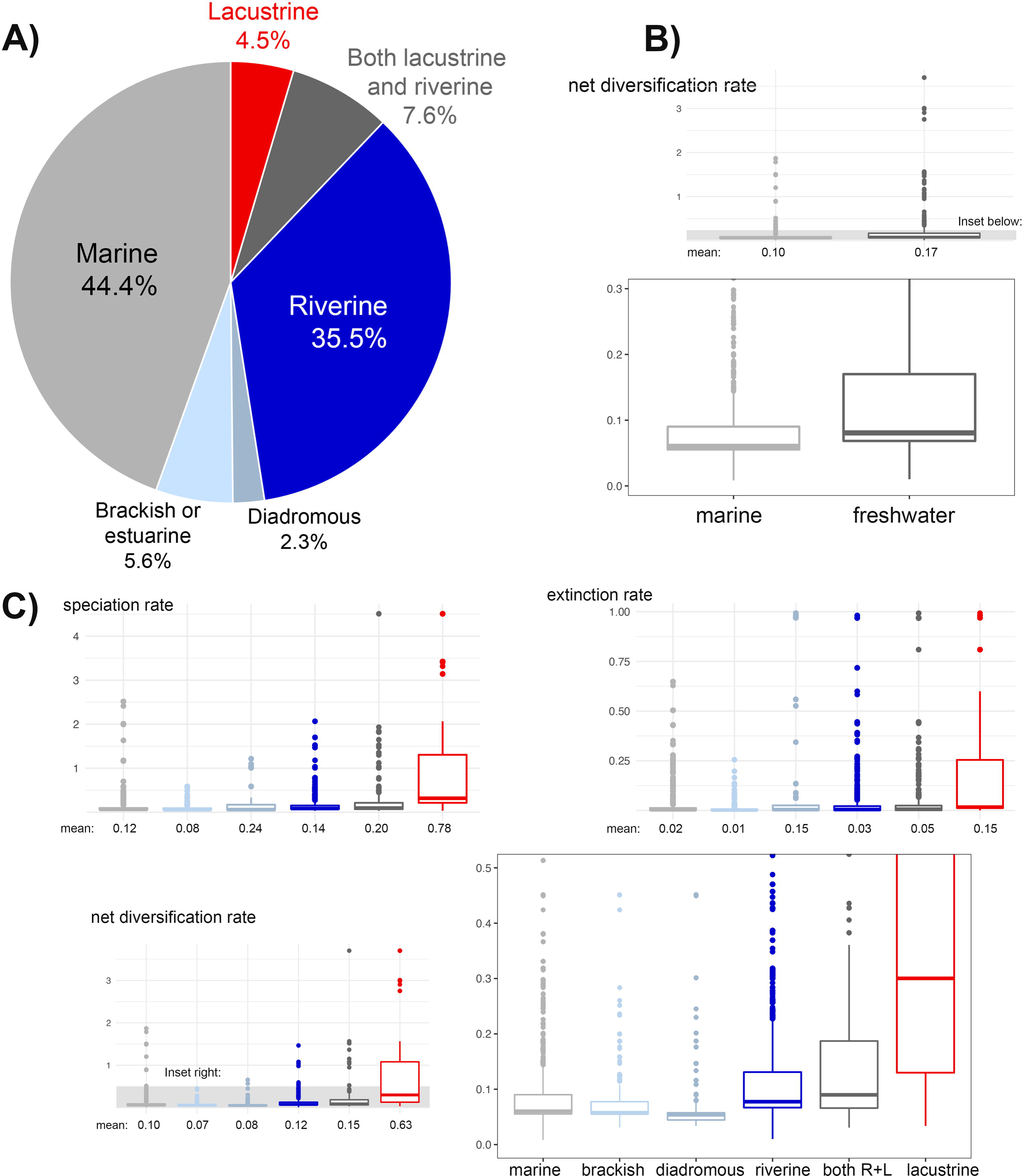
Diversity and diversification rates in aquatic habitats. (A) Proportion of teleost species richness in each habitat (n=11,227 species). (B) Tip-associated net diversification rates in species found exclusively in marine or freshwater habitats (n=10,177 species). (C) Tip-associated speciation, extinction, and net diversification rates among six aquatic habitats. Boxplots summarize rates for individual species, including medians, first and third quartiles, and outliers (outside the 95% confidence interval). Mean rates are printed below each habitat. All rates shown here were calculated using BAMM under a time-varying model of diversification rates (Rabosky et al. 2018; Chang et al. 2019). Results were similar using other methods of calculating diversification rates (Table 3; Table S2–S8).

**Table 1.** Studies comparing diversification rates between (A) freshwater and marine ray-finned fishes, and (B) freshwater, marine, and/or terrestrial habitats using other organisms. Here I note if the study supports the hypothesis that net diversification rates are faster in freshwater than other habitats. See footnotes for method references.

| Study | Clade | Method | Result | Supports hypothesis? |
| --- | --- | --- | --- | --- |
| **A: Ray-finned fishes:** |  |  |  |  |
| Davis et al. 2012 | Terapontidae (grunters) | MEDUSA<sup>1</sup> | Higher freshwater net diversification rates | Yes |
| Carrete Vega and Wiens 2012 | Ray-finned fishes | Method-of-moments<sup>2</sup> | No difference in net diversification rates | No |
| Betancur-R et al. 2012 | Ariidae (sea catfishes) | GeoSSE<sup>3</sup> | No differences in rates | No |
| Bloom et al. 2013 | Atherinopsidae (New World silversides) | BiSSE<sup>4</sup> | Higher freshwater speciation and extinction rates; little difference in net diversification rates | No |
| Santini et al. 2013 | Tetraodontidae (puffers) | MuSSE<sup>5</sup> | Higher marine speciation rates | No |
| Betancur-R et al. 2015 | Ray-finned fishes | GeoSSE | Higher marine net diversification rates | No |
| Guinot and Cavin 2015 | Ray-finned fishes | Fossil family diversity through time | Accumulation of families has slowed through time in marine but not freshwater fishes | Unclear (see text) |
| Tedesco et al. 2017a | Ray-finned fishes | Method-of-moments | Higher freshwater net diversification rates | Yes |
| Miller et al. 2018 | Percomorpha (e.g. cichlids, guppies, reef fishes) | HiSSE<sup>6</sup> | Slightly higher freshwater speciation, extinction and net diversification rates | Yes |
| Rabosky 2020 | Ray-finned fishes | STRAPP<sup>7</sup> | Higher freshwater speciation rates driven by a few clades, but not a general pattern among fishes | Potentially |
| **B: Other organisms:** |  |  |  |  |
| Wiens 2015a | All animal phyla | Method-of-moments | Higher non-marine net diversification rates | Unclear-freshwater and terrestrial combined |
| Wiens 2015b | Vertebrates | Method-of-moments | Higher terrestrial net diversification rates compared to aquatic habitats, but no difference between non-marine and marine habitats | Unclear-freshwater combined with other habitats |
| Poulin 2016 | Fish parasites | Species-to-genus ratios | Higher parasite species-to-genus ratios in freshwater versus marine fishes | Yes |
| Miller and Wiens 2017 | Amniotes | HiSSE | No consistent difference between marine and non-marine rates among clades | Unclear-freshwater and terrestrial combined |
| Kirchhoff et al. 2017 | Myliobatiformes<br>(stingrays) | ClaSSE<sup>8</sup> | No difference in rates between freshwater and<br>marine habitats; brackish rates were lowest | No |
| Nakov et al. 2019 | Diatoms | HiSSE | Higher freshwater speciation, extinction, and<br>net diversification rates | Yes |

1: Alfaro et al. 2009; 2: Magallón and Sanderson 2001; 3: Goldberg et al. 2011; 4: Maddison et al. 2007; 5: Fitzjohn 2012; 6: Beaulieu and O'Meara 2016; 7: Rabosky and Huang 2016; 8: Goldberg and Igić 2012

**Table 2.**
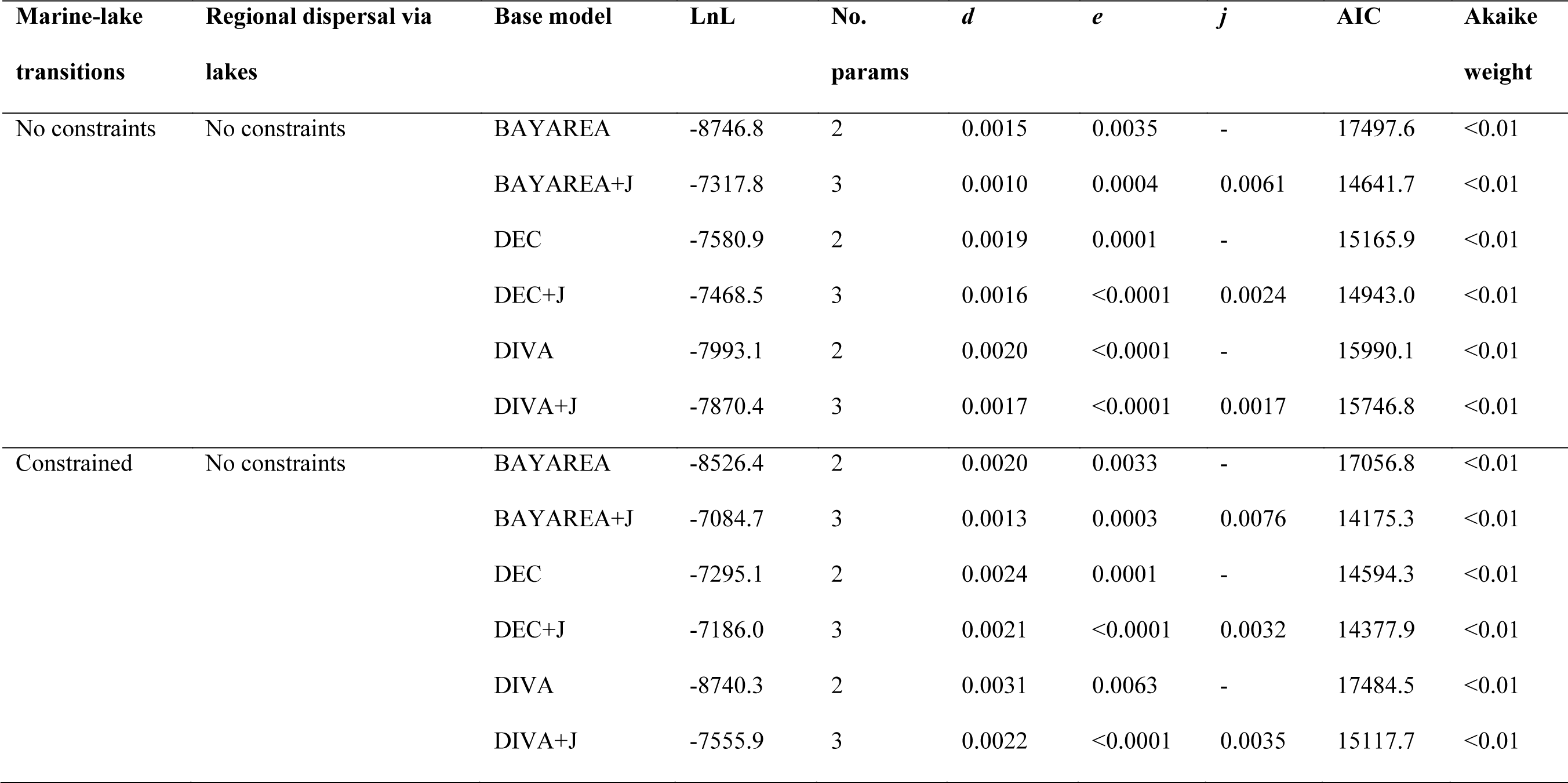
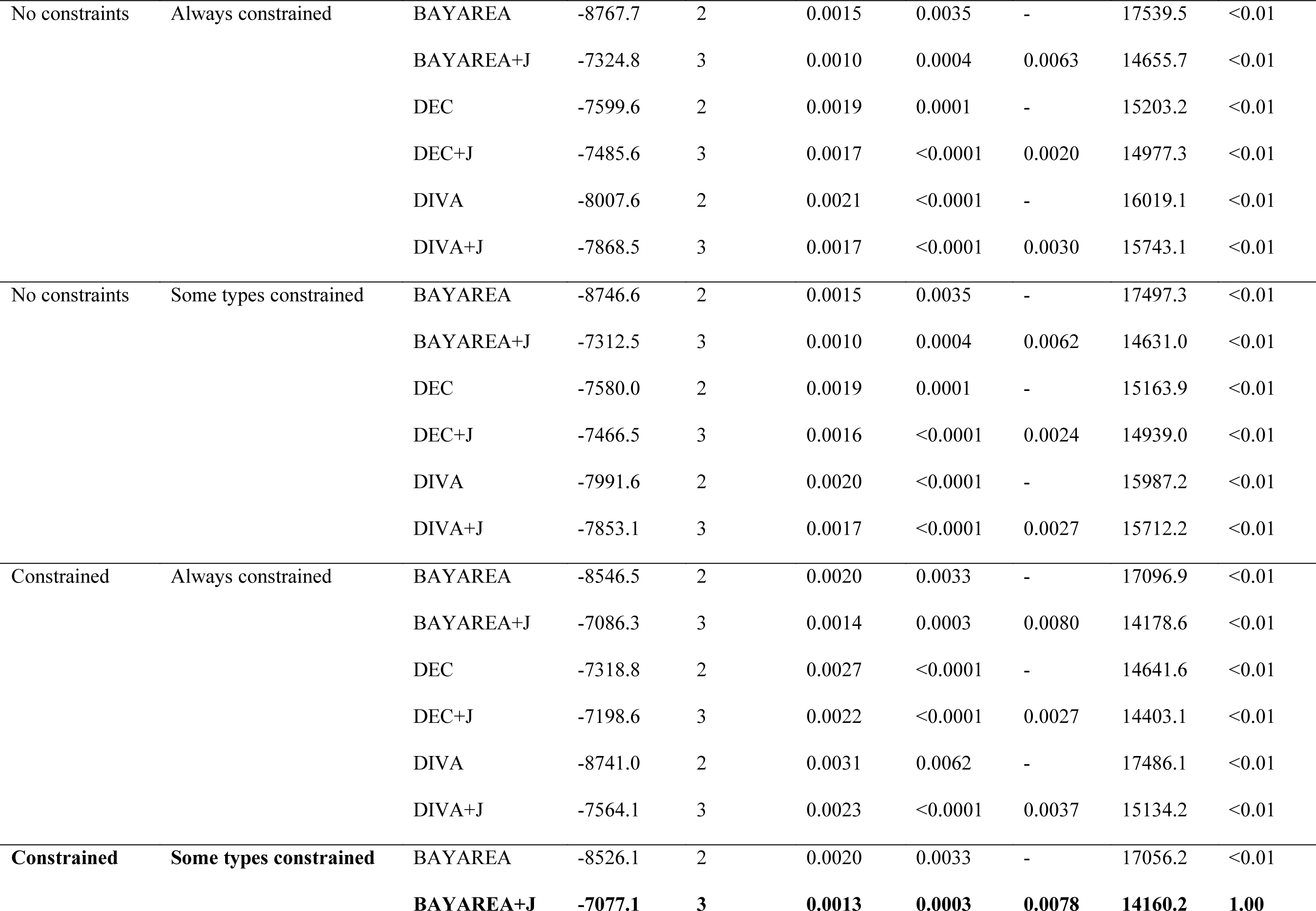
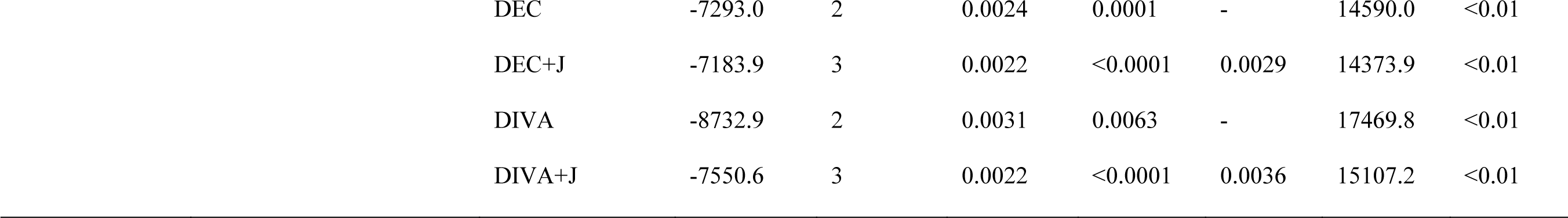
Comparing the fit of alternative biogeographic models fit using BioGeoBEARS (Matzke 2014). Models differed in the constraints set on dispersal in order to test hypotheses about biogeographic connections among lakes. Constraints were set by multiplying the probability of dispersal by 0.05 (more details in Extended Text 2 and Table S11). The best-fitting model (in bold) was used in downstream analyses (Figure 3).

| Marine-lake<br>transitions | Regional dispersal via<br>lakes | Base model | LnL | No.<br>params | *d* | *e* | *j* | AIC | Akaike<br>weight |
| --- | --- | --- | --- | --- | --- | --- | --- | --- | --- |
| No constraints | No constraints | BAYAREA | -8746.8 | 2 | 0.0015 | 0.0035 | - | 17497.6 | <0.01 |
|  |  | BAYAREA+J | -7317.8 | 3 | 0.0010 | 0.0004 | 0.0061 | 14641.7 | <0.01 |
|  |  | DEC | -7580.9 | 2 | 0.0019 | 0.0001 | - | 15165.9 | <0.01 |
|  |  | DEC+J | -7468.5 | 3 | 0.0016 | <0.0001 | 0.0024 | 14943.0 | <0.01 |
|  |  | DIVA | -7993.1 | 2 | 0.0020 | <0.0001 | - | 15990.1 | <0.01 |
|  |  | DIVA+J | -7870.4 | 3 | 0.0017 | <0.0001 | 0.0017 | 15746.8 | <0.01 |
| Constrained | No constraints | BAYAREA | -8526.4 | 2 | 0.0020 | 0.0033 | - | 17056.8 | <0.01 |
|  |  | BAYAREA+J | -7084.7 | 3 | 0.0013 | 0.0003 | 0.0076 | 14175.3 | <0.01 |
|  |  | DEC | -7295.1 | 2 | 0.0024 | 0.0001 | - | 14594.3 | <0.01 |
|  |  | DEC+J | -7186.0 | 3 | 0.0021 | <0.0001 | 0.0032 | 14377.9 | <0.01 |
|  |  | DIVA | -8740.3 | 2 | 0.0031 | 0.0063 | - | 17484.5 | <0.01 |
|  |  | DIVA+J | -7555.9 | 3 | 0.0022 | <0.0001 | 0.0035 | 15117.7 | <0.01 |
| No constraints | Always constrained | BAYAREA | -8767.7 | 2 | 0.0015 | 0.0035 | - | 17539.5 | <0.01 |
|  |  | BAYAREA+J | -7324.8 | 3 | 0.0010 | 0.0004 | 0.0063 | 14655.7 | <0.01 |
|  |  | DEC | -7599.6 | 2 | 0.0019 | 0.0001 | - | 15203.2 | <0.01 |
|  |  | DEC+J | -7485.6 | 3 | 0.0017 | <0.0001 | 0.0020 | 14977.3 | <0.01 |
|  |  | DIVA | -8007.6 | 2 | 0.0021 | <0.0001 | - | 16019.1 | <0.01 |
|  |  | DIVA+J | -7868.5 | 3 | 0.0017 | <0.0001 | 0.0030 | 15743.1 | <0.01 |
| No constraints | Some types constrained | BAYAREA | -8746.6 | 2 | 0.0015 | 0.0035 | - | 17497.3 | <0.01 |
|  |  | BAYAREA+J | -7312.5 | 3 | 0.0010 | 0.0004 | 0.0062 | 14631.0 | <0.01 |
|  |  | DEC | -7580.0 | 2 | 0.0019 | 0.0001 | - | 15163.9 | <0.01 |
|  |  | DEC+J | -7466.5 | 3 | 0.0016 | <0.0001 | 0.0024 | 14939.0 | <0.01 |
|  |  | DIVA | -7991.6 | 2 | 0.0020 | <0.0001 | - | 15987.2 | <0.01 |
|  |  | DIVA+J | -7853.1 | 3 | 0.0017 | <0.0001 | 0.0027 | 15712.2 | <0.01 |
| Constrained | Always constrained | BAYAREA | -8546.5 | 2 | 0.0020 | 0.0033 | - | 17096.9 | <0.01 |
|  |  | BAYAREA+J | -7086.3 | 3 | 0.0014 | 0.0003 | 0.0080 | 14178.6 | <0.01 |
|  |  | DEC | -7318.8 | 2 | 0.0027 | <0.0001 | - | 14641.6 | <0.01 |
|  |  | DEC+J | -7198.6 | 3 | 0.0022 | <0.0001 | 0.0027 | 14403.1 | <0.01 |
|  |  | DIVA | -8741.0 | 2 | 0.0031 | 0.0062 | - | 17486.1 | <0.01 |
|  |  | DIVA+J | -7564.1 | 3 | 0.0023 | <0.0001 | 0.0037 | 15134.2 | <0.01 |
| Constrained | Some types constrained | BAYAREA | -8526.1 | 2 | 0.0020 | 0.0033 | - | 17056.2 | <0.01 |
|  |  | **BAYAREA+J** | **-7077.1** | **3** | **0.0013** | **0.0003** | **0.0078** | **14160.2** | **1.00** |

|  |  |  |  |  |  |  |  |
| --- | --- | --- | --- | --- | --- | --- | --- |
| DEC | -7293.0 | 2 | 0.0024 | 0.0001 | - | 14590.0 | <0.01 |
| DEC+J | -7183.9 | 3 | 0.0022 | <0.0001 | 0.0029 | 14373.9 | <0.01 |
| DIVA | -8732.9 | 2 | 0.0031 | 0.0063 | - | 17469.8 | <0.01 |
| DIVA+J | -7550.6 | 3 | 0.0022 | <0.0001 | 0.0036 | 15107.2 | <0.01 |

**Table 3:**
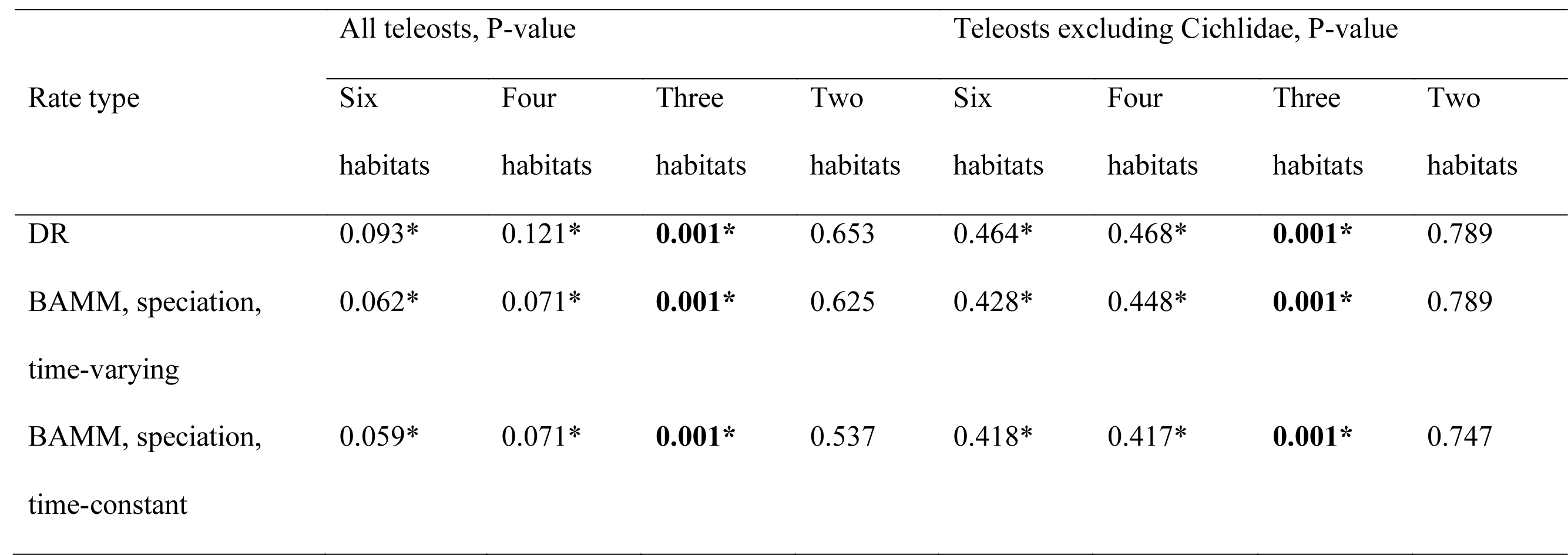
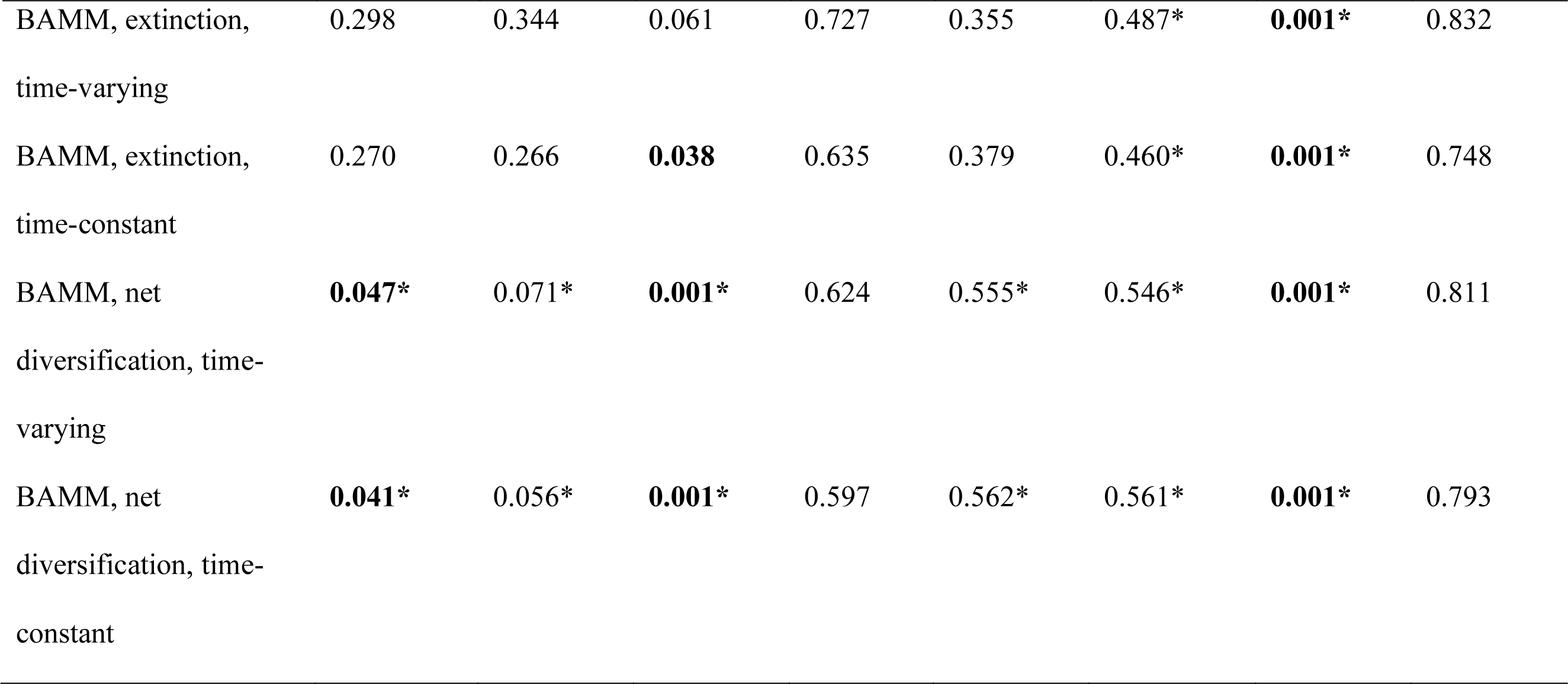
Summary of 56 phylogenetic ANOVA and post hoc tests comparing diversification rates (seven types) among habitat categories (four habitat combinations with and without cichlids). Rates were calculated by Rabosky et al. (2018; see also Chang et al. 2019). Full results including post hoc tests are reported in Tables S2–S8. Six habitats=marine, estuarine, diadromous, riverine, lacustrine, both riverine and lacustrine. Four habitats=marine, riverine, lacustrine, both riverine and lacustrine. Three habitats=riverine, lacustrine, both riverine and lacustrine. Two habitats=marine and freshwater. Boldface=ANOVA was significant. *=Lakes were significantly different from all other habitats in pairwise post hoc tests, even if the ANOVA was not globally significant.

| Rate type | All teleosts, P-value |  |  |  | Teleosts excluding Cichlidae, P-value |  |  |  |
| --- | --- | --- | --- | --- | --- | --- | --- | --- |
|  | Six | Four | Three | Two | Six | Four | Three | Two |
|  | habitats | habitats | habitats | habitats | habitats | habitats | habitats | habitats |
| DR | 0.093* | 0.121* | **0.001\*** | 0.653 | 0.464* | 0.468* | **0.001\*** | 0.789 |
| BAMM, speciation,<br>time-varying | 0.062* | 0.071* | **0.001\*** | 0.625 | 0.428* | 0.448* | **0.001\*** | 0.789 |
| BAMM, speciation,<br>time-constant | 0.059* | 0.071* | **0.001\*** | 0.537 | 0.418* | 0.417* | **0.001\*** | 0.747 |
| BAMM, extinction,<br>time-varying | 0.298 | 0.344 | 0.061 | 0.727 | 0.355 | 0.487* | **0.001\*** | 0.832 |
| BAMM, extinction,<br>time-constant | 0.270 | 0.266 | **0.038** | 0.635 | 0.379 | 0.460* | **0.001\*** | 0.748 |
| BAMM, net<br>diversification, time-varying | **0.047\*** | 0.071* | **0.001\*** | 0.624 | 0.555* | 0.546* | **0.001\*** | 0.811 |
| BAMM, net<br>diversification, time-constant | **0.041\*** | 0.056* | **0.001\*** | 0.597 | 0.562* | 0.561* | **0.001\*** | 0.793 |

Both ‘freshwater’ and ‘marine’ are broad categories that encompass diverse habitats and lineages. Diversification rates are known to vary among marine fishes: for example, diversification is faster on coral reefs than in other marine habitats (Santini et al. 2013; Tedesco et al. 2017a). I am not aware of similar studies in fishes comparing diversification rates among different freshwater habitats. An examination of diversification-rate variation among freshwater fish clades (Rabosky et al. 2013; Seehausen and Wagner 2014) shows that some species-rich clades do not have fast rates (e.g. cyprinids), while some groups with fast rates are not species-rich (e.g. sticklebacks). To what extent can these patterns be explained by differences in habitat? The most commonly proposed mechanism for faster freshwater diversification rates, that freshwater habitats give more opportunities for population subdivision than marine habitats, implies that allopatric speciation has generated much of the world’s freshwater diversity (Lynch 1989). This may be the case, as major river systems such as the Amazon are among the most diverse freshwater habitats (Lévêque et al. 2008; Albert et al. 2020; Miller and Román-Palacios 2021). The fragmentation of rivers has been positively related to endemism and cladogenesis in fishes (Dias et al. 2013). However, lakes are also known for their diverse faunas (Lévêque et al. 2008). Ancient lakes such as the African rift lakes and Lake Baikal contain species flocks, or monophyletic groups that underwent rapid speciation within a single lake (Cohen 1995; Cristescu et al. 2010; Wagner et al. 2012; Seehausen 2015). While allopatric speciation is also common within and among lakes (Parenti 1984; Seehausen and van Alphen 1999; Rico and Turner 2002; Hrbek et al. 2002; Martin et al. 2015), intralacustrine radiations represent some of the best-supported cases of sympatric speciation across the Tree of Life (Schluter and McPhail 1993; Bolnick and Fitzpatrick 2007; Seehausen and Wagner 2014; Kautt et al. 2020). The relative contribution of speciation in rivers versus lakes to freshwater diversity overall is an open question (Seehausen and Wagner 2014).

A related question is the pace of speciation inherent to allopatric versus sympatric speciation modes. Speciation is thought to proceed slowly in allopatry because genetic differences between isolated populations accumulate passively (genetic drift) or through gradual local adaptation. Reproductive isolation may be achieved faster in sympatry than in allopatry because disruptive selection may actively select against hybridization (Bush 1975; McCune and Lovejoy 1998; Crow et al. 2010; Germain et al. 2020). McCune and Lovejoy (1998) showed that sympatric lacustrine sister-species had fewer genetic differences between them than allopatric riverine sister-species, suggesting that reproductive isolation was achieved faster between sympatric sister-species. However, later studies in other fishes found that speciation can also be rapid in allopatry in both lacustrine and riverine habitats (Hrbek et al. 2002; Near and Bernard 2004). It is unclear what generalities can be drawn from these studies limited to a few well-studied fish clades.

There are at least two reasons why we may expect differences in net diversification rates in fishes found in riverine versus lacustrine habitats. First, these habitats may be differently conducive to alternative speciation modes which differ in their time to completion. Allopatric speciation seems to be the dominant mode in rivers because rivers become fragmented and change course through time. Evidence for this statement comes from the allopatric distribution of closely related species in river systems around the world (Near and Bernard 2004; Dias et al. 2013; Albert et al. 2020; Sholihah et al. 2021). A few cases of sympatric or parapatric speciation within rivers are well-supported (mormyrid fishes, Sullivan et al. 2002; African *Teleogramma* cichlids, Alter et al. 2017; Neotropical *Crenicichla* cichlids, Burress et al. 2018), but in general sympatric speciation appears to be less common in rivers than in lakes (Seehausen and Wagner 2014; Seehausen 2015). If speciation rates are faster in sympatry than in allopatry, then we may expect lacustrine fishes to have faster speciation and net diversification rates than riverine fishes (McCune and Lovejoy 1998).

Second, rivers and lakes may differ in ecological opportunity (Yoder et al. 2010). Early literature posited that lakes should have faster speciation rates for ecological and structural reasons. Briggs (1966) and Lowe-McConnell (1969) thought that rates of speciation should generally be faster in peripheral habitats than in centers of species diversity, with both authors comparing rivers and lakes. Lowe-McConnell (1969) and later Seehausen (2015) also argued that there may be fewer available niches in rivers: since rivers are locally unstable, riverine fishes are more likely to be opportunistic feeders. The more stable conditions of lakes allow for trophic specialization. In addition, since lakes tend to be wider and deeper than most rivers, they offer three dimensions to partition habitats: onshore-offshore, shallow-deep, and along the shoreline (Seehausen 2015). This dimensionality may encourage divergent selection, promote species coexistence and allow for faster speciation (Germain et al. 2020).

Differences in diversification rates among freshwater habitats may be a crucial missing piece to interpreting comparisons between freshwater and marine diversification. Specifically, an alternative hypothesis to explain the freshwater paradox is that freshwater diversity generally accumulated over long periods of time rather than at a very fast rate (Wiens 2012; Carrete Vega and Wiens 2012; Miller and Román-Palacios 2021). This could still be true if most freshwater species arose through fine-scale allopatric speciation as predicted (e.g., Grosberg et al. 2012). If so, the higher diversification rates sometimes observed in freshwater relative to marine fishes (Table 1) may actually be driven by rapid speciation within lakes, not allopatric speciation in rivers. This would call into question the most commonly proposed mechanism for why diversification rates seem to differ between biomes.

In this study, I compared rates of speciation, extinction, and net diversification between marine and freshwater teleost fishes. However, unlike past studies, I compared these rates in riverine and lacustrine habitats separately rather than treating freshwater as a single category. I found that diversification rates in riverine and marine fishes are actually similar on average. However, diversification rates in lakes are much faster than in other aquatic habitats. These results show that more nuance is needed when comparing marine and freshwater diversification patterns.

## MATERIAL AND METHODS

### Data acquisition

In this study I used the largest available time-calibrated phylogeny of ray-finned fishes for all analyses (Rabosky et al. 2018). This maximum likelihood phylogeny was constructed by aggregating genetic data from many past studies, and contains 11,638 species (36.9% of described actinopterygian fishes). I limited the present study to teleost fishes, a group containing 99.8% of ray-finned fishes, because living non-teleost clades are much older with many extinct members (Betancur-R et al. 2015) and it may not be appropriate to compare their diversification rates with teleosts. I removed 51 tips that were duplicate or unresolved species and 48 tips belonging to the four living non-teleost orders (Polypteriformes, Acipenseriformes, Lepisosteiformes, and Amiiformes), leaving phylogenetic data for 11,539 species.

To assign each species in the phylogeny to aquatic habitat categories I used descriptions of geographic range and habitat in FishBase (Froese and Pauley 2019), the IUCN Red List version 6.2 (2020), and a recent compilation of habitat in fishes (Corush 2019). I identified species that were marine, freshwater, diadromous, or estuarine/brackish. Freshwater species were further categorized as riverine, lacustrine, or occurring in both lakes and rivers. Some species were found in additional freshwater habitats including swamps, ponds, temporary pools, caves, and springs. I found that diversification rates in these habitats were similar to those in rivers (details in Extended Text 1; Figure S1), perhaps because these habitats tend to be created or maintained by fluvial action. Species found in these other habitats were simultaneously found in rivers much more often than in lakes (Table S1). For these reasons, I used the species’ distribution in rivers and/or lakes as the sole basis for its habitat categorization. This approach was not applicable to the (very few) species that were endemic to these other habitats (Table S1). Rather than exclude these species or include many additional habitat categories in my analyses, I coded these species as riverine.

Of the 11,539 species in the phylogeny, I identified 4,984 as exclusively marine (44.4%), 631 as brackish, estuarine or euryhaline (5.6%), 263 as diadromous (2.3%), 3,985 as riverine (35.5%), 510 as lacustrine (4.5%), and 854 in both rivers and lakes (7.6%). I removed the remaining 312 species with unclear habitat affinities from analyses, leaving 11,227 species.

### Comparing diversification rates among habitats

I compared diversification rates among habitat types using tip-associated rates (Title and Rabosky 2019) calculated for each species by Rabosky et al. (2018). Rates associated with each species were downloaded using the R package fishtree version 0.2.0 (Chang et al. 2019). I compared seven alternative rate types: (1) the DR statistic (Jetz et al. 2012) which was calculated from the phylogeny of ray-finned fishes with all missing species imputed on the phylogeny, (2) speciation calculated by Bayesian Analysis of Macroevolutionary Mixtures (BAMM; Rabosky 2014) under a time-varying model on the phylogeny with genetic data only, (3) speciation calculated by BAMM under a time-constant model, (4) extinction calculated by BAMM under a time-varying model, (5) extinction calculated under a time-constant model, (6) BAMM net diversification rates (speciation minus extinction rates) under a time-variable model, and (7) BAMM net diversification rates under a time-constant model. Note that tip rates were not available for 172 species (Chang et al. 2019). Analysis settings used to obtain these rates are detailed in the original study (Rabosky et al. 2018). The DR statistic for any given species is estimated from the number of branches (i.e. splitting events) between the species and the root of the tree and the length of those branches (Jetz et al. 2012). This type of rate estimate resembles speciation rates more closely than net diversification rates (Title and Rabosky 2019). BAMM uses a model of diversification to simulate a posterior distribution of rate shift configurations, in which taxa in the same rate shift cohort have identical tip rates. While -SSE models can also be used to compare diversification rates among habitats (Table 1), I chose to use rates calculated using methods agnostic to traits (DR, BAMM). This is because it is straightforward to compare rates among several habitat categories at once using these trait-agnostic rates, while -SSE models become very computationally intensive when more than two traits are considered especially at this broad phylogenetic scale.

I statistically compared the mean tip rates of habitats using phylogenetic ANOVA (Garland et al. 1993) implemented in phytools version 0.6–44 (Revell 2012). The *phylANOVA* function in phytools implements a simulation-based post hoc test to compare pairwise differences in means, and also uses a Holm-Bonferroni correction for multiple comparisons. I made a total of four habitat comparisons with and without cichlids, using each of the seven rate types (see above), for a total of 56 comparisons. First, I compared tip-associated diversification rates between species in two habitats, as classically done (Table 1): exclusively marine and exclusively freshwater (excluding diadromous or estuarine species). Second, I compared rates among six habitat categories: riverine, lacustrine, both riverine and lacustrine, marine, estuarine, and diadromous. Third, I compared four habitats: exclusively marine, riverine, lacustrine, and both riverine and lacustrine. I excluded diadromous and euryhaline species from this comparison because the habitat where speciation occurred may be unclear when species interact with both freshwater and marine habitats. Fourth, I compared only freshwater habitat types (three categories: riverine, both riverine and lacustrine, and lacustrine). Finally, I repeated these four habitat comparisons after removing the family Cichlidae. This was to test if cichlids were driving any differences in rates observed among habitats. Lacustrine cichlids possess the fastest diversification rates among fishes (Burress and Tan 2017; McGee et al. 2020) and so may have an overwhelming influence on the results.

### Alternative divergence times

The age of clades can potentially affect diversification rate estimates by shortening or lengthening branch lengths within the clade. Several new divergence time estimates are available for fishes using phylogenomic techniques. However, the dense sampling of the Rabosky et al. (2018) phylogeny is needed to estimate tip-associated diversification rates. To test whether alternative divergence estimates have cascading effects on diversification rate differences among habitats, I used the “congruification” approach of Eastman et al. (2013). This approach uses a reference phylogeny, which is a tree containing few exemplar tips representing higher taxa, to time-calibrate a target phylogeny with shared higher taxa but denser species-level sampling. Thus, the congruification approach attempts to join the efforts of systematists using alternative approaches to build and date trees (few exemplar taxa with many base pairs per species versus broad sampling with less data per species).

I used the genomic phylogenies of Hughes et al. (2018) for teleosts and Alfaro et al. (2018) for Acanthomorpha (spiny-rayed fishes) as alternative target phylogenies. Notably, Alfaro et al. (2018) found that most major clades within acanthomorphs diverged near the K-Pg boundary (66 mya), more recently than in Hughes et al. (2018) or Rabosky et al. (2018). I downloaded the undated maximum likelihood phylogeny of fishes (branch lengths scaled by molecular substitutions) from the Fish Tree of Life website (https://fishtreeoflife.org/downloads/). I implemented the congruification using the function *congruify.phylo* in the R package geiger version 2.0.7 (Pennell et al. 2014) which uses TreePL (Smith & O’Meara 2012) to time-calibrate the tree. This procedure resulted in two rescaled trees: a phylogeny of teleosts with identical sampling as Rabosky et al. (2018) but dates similar to Hughes et al. (2018), and a phylogeny of acanthomorphs with identical sampling as Rabosky et al. (2018) but dates similar to Alfaro et al. (2018).

I calculated the DR statistic using each rescaled tree and compared diversification rates among habitat categories as detailed above. This set of analyses added 16 comparisons (4 habitat combinations, with and without cichlids, using each re-scaled phylogeny) to the 56 performed using the original phylogeny for a total of 72 comparisons among three phylogenies.

### Biogeographic model fitting

I found that global freshwater biodiversity was mostly found in rivers, with much lower richness distributed among lakes (Fig. 1). Species richness among habitats can be explained by three processes: in-situ speciation, extinction, and dispersal (transitions) among habitats (Ricklefs 1987; Wiens 2012). To understand why there are much fewer species in lakes than in rivers, I used biogeographic models to trace the history in each habitat along the phylogeny (Rabosky et al. 2018). I fit alternative biogeographic models implemented in the program BioGeoBEARS version 1.1.2 (Matzke 2014). I summarize my approach to model fitting here and give more details in Extended Text 2. Note that these analyses are similar to those in Miller and Román-Palacios (2021). The goal of that study was to compare colonization patterns and diversification rates among the freshwater faunas of biogeographic regions, but did not distinguish between lakes and rivers within these regions.

I modeled dispersal among 13 habitat-region combinations. I included six biogeographic regions following Leroy et al. (2019), who used clustering algorithms to identify a bioregionalization scheme based on shared freshwater fish diversity (region names also follow this study). The six regions were: (1) Nearctic, (2) Neotropical, (3) Palearctic, (4) Ethiopian, (5) Sino-Oriental, and (6) Australia. Note that Miller and Román-Palacios (2021) used the earlier regionalization of Tedesco et al. (2017b) which has minor differences in boundaries. Species in these six regions were further assigned to riverine and/or lacustrine habitats, giving 12 region-habitat combinations (e.g. Neotropical rivers and Neotropical lakes). Marine and estuarine species were coded as belonging to a 13^th^ “marine” region. I restricted the root of the teleost phylogeny to the marine state in accordance with the fossil record of fishes (Betancur-R et al. 2015). Following prior biogeographic studies in freshwater systems (Toussaint et al. 2017; Miller and Román-Palacios 2021), I restricted dispersal probabilities in accordance with changing regional connectivity through time (Table S11). These dispersal constraints were common to all models.

I also compared the fit of six model variants with modified transition matrices in order to test hypotheses about the role of lakes in dispersal. In variant 1, dispersal among biogeographic regions was unrelated to habitat. For example, dispersal from the Neotropical to the Nearctic region could occur as easily within the riverine as the lacustrine state, and marine-to-lacustrine transitions could occur as easily as marine-to-riverine transitions. In variant 2, the probability of transition between marine and lacustrine habitats was reduced (multiplied by a scalar value of 0.05). This model would be preferred if most transitions from marine to freshwater habitats occurred via rivers. In variant 3, the probability of dispersal among biogeographic regions was much lower via lakes than via rivers (i.e., all regional dispersal probabilities within the lacustrine state were multiplied by 0.05). For example, dispersal from Neotropical to Nearctic rivers was more likely than dispersal from Neotropical to Nearctic lakes. This model would be preferred if most biogeographic dispersal occurred within rivers. In variant 4, restrictions on lacustrine dispersal were relaxed such that lacustrine dispersal occurred at the same probability as riverine dispersal across adjacent regions or short marine barriers, but lacustrine dispersal was downweighed across large overland or overseas barriers (multiplied by 0.05). For example, dispersal from Neotropical to Nearctic rivers was equally likely as from Neotropical to Nearctic lakes; however, dispersal from the Neotropical rivers to Ethiopian rivers was more likely than the equivalent through lakes. Variant 5 combined marine-lake constraints with strict constraints on biogeographic dispersal through lakes. Variant 6 combined marine-lake constraints with relaxed constraints on dispersal through lakes.

In total, I fit 36 alternative biogeographic models (Table 2). These were six models with varying dispersal matrices for each of six base model types (DEC, DEC+J, DIVALIKE, DIVALIKE+J, BAYAREALIKE, and BAYAREALIKE+J). Models with +J denote the addition of a jump parameter allowing cladogenetic dispersal; otherwise, an assumption of models is that all dispersal occurs anagenetically (further differences among models are described in Matzke 2014). The best-fit model was chosen by comparing Akaike weights (Burnham and Anderson 2002).

### Biogeographic patterns

In order to identify independent dispersal events and visualize uncertainty in the ancestral reconstruction, I simulated 100 biogeographic stochastic maps (Dupin et al. 2017) using the best-fit model. Each individual simulation is a realized history that is possible given the model and data, including the time and location on the branches for biogeographic events. Averaging over all of these simulations will approximate the ancestral state probabilities calculated by the model.

To understand why there are fewer species in lakes than rivers, I used biogeographic stochastic maps to answer the following questions. First, what is the rate of transition into and out of lakes relative to other habitats? To obtain habitat transition rates, I counted the number of transitions to marine, riverine only, lacustrine only, and riverine+lacustrine states (combining these counts among regions for each habitat type). The count of events alone can be a misleading measure of the transition rate. As an example, if most freshwater lineages occur in rivers then there may be more transitions from rivers to lakes simply by chance. To convert these counts into a transition rate, for each transition type I divided the number of transitions from the source habitat to the receiving habitat by the sum of branch lengths reconstructed in the source habitat. For example, to calculate the transition rate from marine to riverine I divided the count of these transitions by the sum of branch lengths reconstructed in the marine state. These counts vary slightly among stochastic maps, so I did this for all 100 simulations individually and then took the mean of the 100 rate estimates.

Second, what is the contribution of individual transition events to overall freshwater diversity, and are there replicated patterns shared by habitats across biogeographic regions? More specifically, for each lineage representing an independent transition to a habitat I counted the number of descendants of that transition (species richness; note that this count only included species remaining in that habitat by the present), the mean tip-associated diversification rate among the descendants (DR), and the age of the lineage (time of the transition). Then, I took the mean of these values among lineages found in each habitat and biogeographic region. Again, the estimated location of habitat transitions on the phylogeny varies slightly among stochastic maps, so I did this for all 100 simulations. Note that the richness of independent habitat transitions was inferred by counting tips in the phylogeny descended from these transitions, because I could not assign unsampled species in the phylogeny to individual biogeographic events. I assessed the potential sensitivity of these results to biased sampling in Extended Text 3.

Finally, I visualized the accumulation of diversity through time in each freshwater habitat and region using the lineage-through-space-and-time approach of Skeels (2019) implemented using the R package ltstR version 0.1.0. This approach is analogous to a lineage-through-time plot but uses the output of biogeographic stochastic mapping to separate temporal patterns of species accumulation by region.

## RESULTS

### Diversification rates in aquatic habitats

The results of 56 rate-habitat comparisons using the phylogeny of Rabosky et al. (2018) are summarized in Table 3 and given in full in Tables S2–S8. When comparing only two habitat categories, net diversification rates in marine and freshwater teleosts were broadly overlapping but slightly higher in freshwater (mean net diversification rate=0.10 species/million years in marine and 0.17 in freshwater under a BAMM time-varying model; Fig. 1). This difference was not significant using phylogenetic ANOVA no matter whether speciation, extinction or net diversification rates were used, the method to calculate these rates, or whether the family Cichlidae was included (Table 3).

When comparing six habitat categories, lacustrine fishes had much faster speciation, extinction and net diversification rates than other aquatic habitats (mean net diversification rate=0.63 under a BAMM time-varying model; Fig. 1). Surprisingly, rates in marine and riverine habitats were similar (mean net rate=0.10 in marine and 0.12 in riverine habitats). Species found in both rivers and lakes diversified at rates more similar to rivers than to lakes (mean net rate=0.15). Estuarine and diadromous species had slightly lower net diversification rates than other habitats (mean net rate=0.07 in estuaries and 0.08 in diadromous species).

Comparisons of net diversification rates among six habitat categories using phylogenetic ANOVA were significant when including all teleosts (BAMM time-varying model: P=0.047; BAMM time-constant model: P=0.041). Differences in speciation and extinction rates were not significant, nor were net diversification rates after excluding Cichlidae (Table 3). Still, lakes were significantly different from the other five habitats in speciation and net diversification rates in all pairwise post hoc tests, even when the ANOVAs were not globally significant. This was true using all methods to calculate rates and whether or not cichlids were included (Table 3). Comparing four habitat categories (excluding estuarine and diadromous species) resulted in the same pattern as comparing six: phylogenetic ANOVAs were not globally significant, but lakes were significantly different in speciation and net diversification rates from the other three habitats in pairwise post hoc tests. When comparing just the three freshwater habitats (riverine, lacustrine, and both riverine and lacustrine), phylogenetic ANOVAs became globally significant in speciation, extinction and net diversification rates. This was driven by faster rates in lakes compared to rivers or rivers+lakes, no matter the rate method or exclusion of cichlids. Among all 56 comparisons, pairwise tests were never significant between other habitat categories except for one other case (diadromous species had faster extinction rates than estuarine or marine species; Tables S3, S6).

Taken together, these results suggest that: (1) lakes have faster speciation and net diversification rates than other freshwater habitats (and potentially faster extinction, though this was not significant), (2) rapid lacustrine diversification rates are not limited to cichlids, and (3) marine rates are similar to riverine rates. As a consequence of the latter result, comparing just two categories (freshwater and marine) can mask the signal of rapid lacustrine diversification since riverine species are much more common than lacustrine species (Fig. 1). The similarity between marine and riverine diversification rates also explains why phylogenetic ANOVAs were not significant when comparing four categories but were significant when comparing the three freshwater categories (Table 3).

These patterns held when using phylogenies re-dated in reference to phylogenomic hypotheses (Alfaro et al. 2018; Hughes et al. 2018). The results of the 16 additional comparisons using re-dated trees are summarized in Fig. 2 and shown in full in Tables S9–S10. Unsurprisingly, re-scaling the phylogeny of Acanthomorpha using the Alfaro et al. (2018) hypothesis resulted in slightly faster rates for all habitats because the younger divergence time estimates shortened branch lengths across the entire phylogeny. Still, diversification rates (DR statistic) were generally similar before and after time-calibrating the Rabosky et al. (2018) tree based on either genomic phylogeny (Fig. 2; Fig. S2). Lacustrine species still had the fastest diversification rates among aquatic habitats as shown by pairwise post hoc tests.

**Figure 2.**
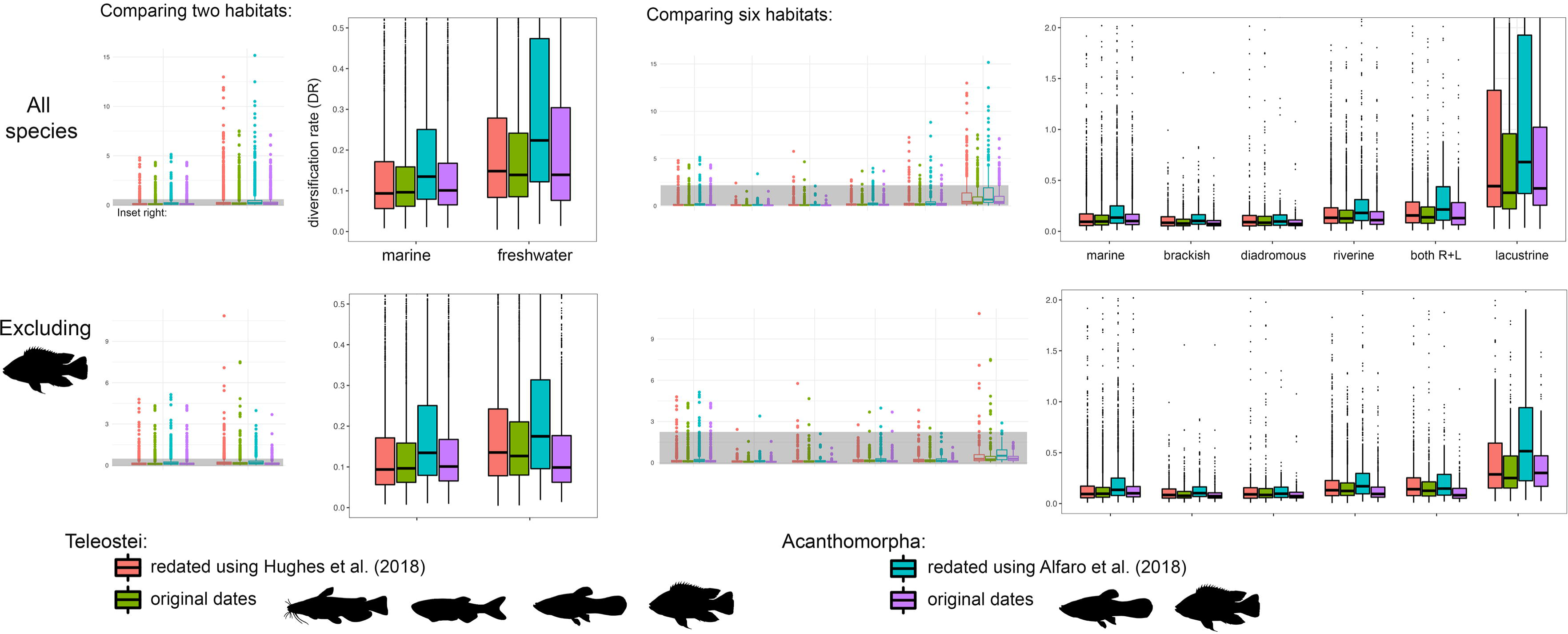
Diversification rate variation by habitat using alternative divergence time estimates and clade selection. All rates shown were calculated using the DR statistic (full results in Tables S2–S10). Boxplots summarize rates for individual species, including medians, first and third quartiles, and outliers (outside the 95% confidence interval). Left panels compare rates in species exclusive to marine and freshwater habitats; right panels compare rates among six aquatic habitats. Top panels compare rates using all members of the clade; right panels compare rates after excluding the family Cichlidae. For teleosts, red bars represent rates calculated from the Rabosky et al. (2018) phylogeny redated using Hughes et al. (2018) as a reference; green bars represent rates using the original tree. For acanthomorphs (spiny-rayed fishes), blue bars represent the Rabosky et al. (2018) phylogeny redated using Alfaro et al. (2018) as a reference; purple bars represent rates using the original tree (Acanthomorpha alone). Fish icons from PhyloPic (http://phylopic.org/) with credit to Lily Hughes, Tyler McCraney, and Milton Tan.

### Habitat transitions and the formation of freshwater diversity

The best-fitting biogeographic model (BAYAREA+J with dispersal matrix variant 6; Table 2) constrained marine-lacustrine transitions to be less probable than marine-riverine transitions. This model also applied some constraints on regional dispersal while in the lacustrine state. Rates of cladogenetic (“jump”) dispersal were six times higher than rates of anagenetic dispersal (*j*=0.0078 events/million years versus *d*=0.0013; Table 2), suggesting that founder events are an important means of dispersal and speciation among freshwater habitats and regions.

In general, marine and riverine habitats were reconstructed in deep ancestral nodes, while the lacustrine state was concentrated towards more recent nodes (Fig. 3A). The rate of transition from marine to riverine habitats was seven times higher than the rate from marine to lacustrine (0.0007 versus 0.0001 events per million years; Fig. 3B). The transition rate from the joint river+lake state into lakes alone was over twice as high as the rate of direct dispersal from rivers to lakes (0.0031 versus 0.0013). Thus, the most typical means of becoming a lacustrine fish was to first transition from marine to riverine habitats, then from rivers alone to both rivers and lakes, then finally becoming endemic to lakes.

**Figure 3.**
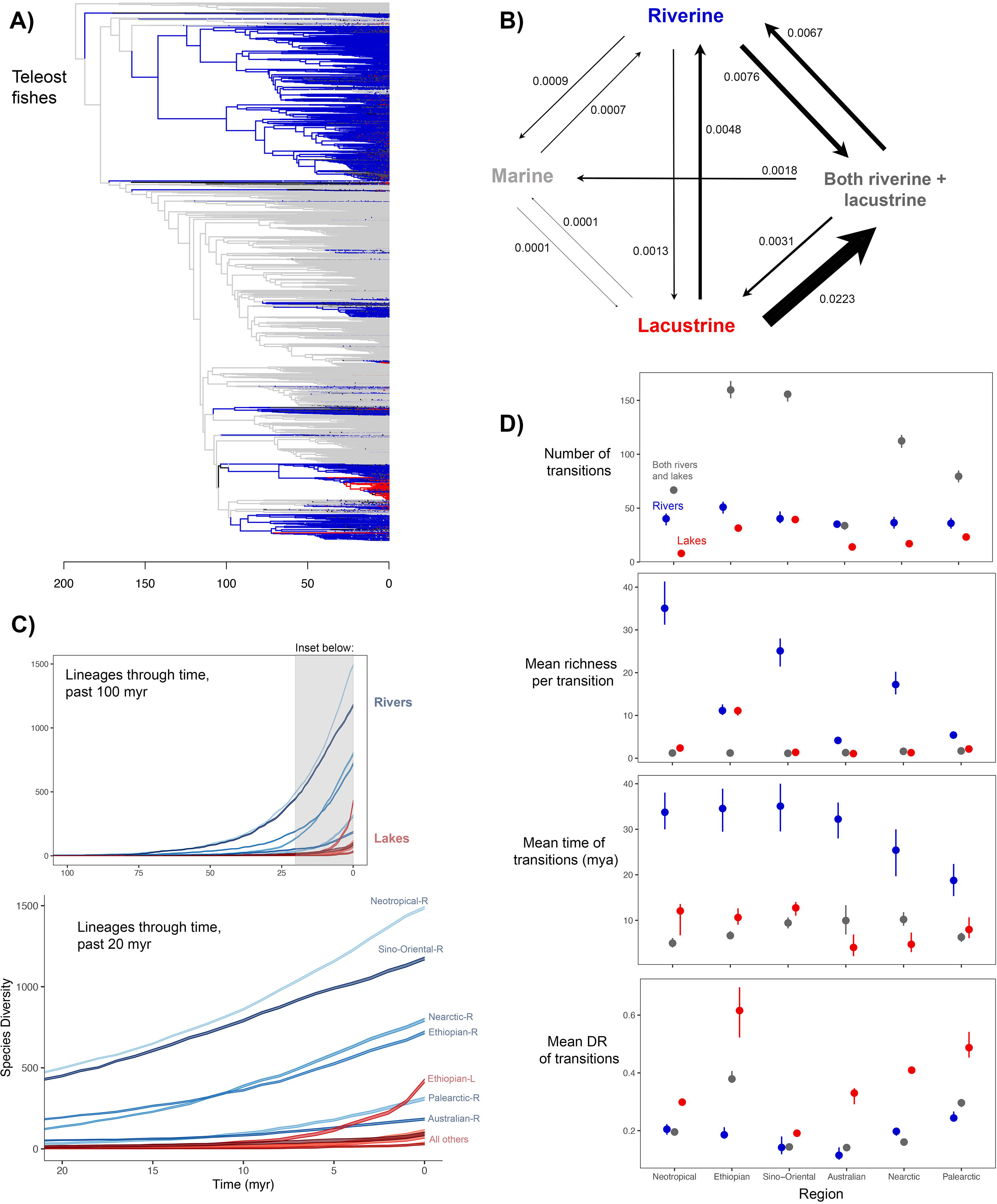
Results of biogeographic and habitat modelling. In panels A and B, biogeographic regions were combined to illustrate habitat transitions overall. In C and D, habitats are shown separately for each region. (A) Ancestral habitats based on best-fit model (Table 2). Here, light grey=marine, blue=riverine, red=lacustrine, dark grey=both riverine and lacustrine, and black=uncertain (no habitat with probability >80%). (B) Transition rates among habitats. Black arrows are sized to scale of transition rates. Number is the mean transition rate calculated from 100 biogeographic stochastic maps. (C) Lineages-through-space-and-time plot (Skeels 2019) of total riverine and lacustrine diversity within six biogeographic regions. The width of lines represents the range of values observed among 100 stochastic maps. Bioregionalization follows Leroy et al. (2019). (D) Properties of lineages representing independent habitat transitions by region. From top to bottom: number of independent transitions into that habitat, mean number of species descending from transitions, mean time of transitions, and mean tip-associated diversification rate (DR) of descendants. Points show the mean and range of mean values among 100 stochastic maps.

Transition rates out of lakes were higher than transition rates into lakes (Fig. 3B). The rate of transition from lakes alone back to rivers+lakes was about seven times higher than the reverse (0.0223 events/million years versus 0.0031). The direct dispersal rate from lakes to rivers was over three times as high as the reverse (0.0048 versus 0.0013). This suggests that of the few lineages that become endemic to lakes, many will eventually enter rivers again.

I compared four metrics characterizing lineages associated with habitat transitions: the number of independent habitat transitions, mean richness per habitat transition, mean timing of habitat transitions, and mean diversification rate (DR). Differences in these metrics among rivers, lakes, and rivers+lakes were generally shared across the six biogeographic regions (Fig. 3D). Transitions into the rivers+lakes state were much more common than transitions into rivers or lakes alone for four of six regions (all but Neotropical and Australian regions). Transitions into lakes were the least common in five of six regions (all but Sino-Oriental).

Mean species richness per habitat transition was highest for the riverine state in five of six regions (Fig. 3D). The mean richness of transitions into lakes was low in all but the Ethiopian region, where richness-per-transition was similar between rivers and lakes. Note that these richness differences cannot be attributed to sampling biases by habitat: lacustrine species were better sampled in the phylogeny than riverine species (Extended Text 3). In all six regions, lineages transitioning into the rivers+lakes state were usually limited to single species, suggesting that this habitat is a transitional state between rivers and lakes.

In all six regions, riverine transitions were older on average than transitions into the rivers+lakes or lacustrine states (Fig. 3D).

The mean diversification rate (DR) of lacustrine lineages was higher than that of the other habitat states in all regions. The difference between lacustrine and riverine diversification rates was large in the Ethiopian, Australian, Nearctic and Palearctic regions, and smaller in the Neotropical and Sino-Oriental regions (Fig. 3D).

The temporal patterns of lineage accumulation in rivers and lakes worldwide reflect these differences in transition rates, transition timing, and diversification rates with habitat (Fig. 3C). All six regions have much greater diversity contained in rivers than lakes, even though average diversification rates were always higher in lakes (Fig. 3C). Freshwater diversity has been highest overall in Neotropical and Sino-Oriental rivers for the past 75 million years, reinforcing the role of time for explaining these mega-diverse faunas (Miller and Román-Palacios 2021). Only the Ethiopian region has lacustrine diversity rivalling total riverine diversity in some other regions, which is due to the exceptionally fast diversification rates in African lakes (Fig. 3D). Note that time and diversification-rate patterns characterizing biogeographic regions themselves were consistent with those found by Miller and Román-Palacios (2021).

Taken together, these biogeographic results (Table 2; Figure 3) suggest that most global freshwater diversity is riverine in spite of faster lacustrine diversification rates for the following reasons. First, most lineages transitioning to freshwater from marine biomes enter rivers first. Second, transitions from rivers to lakes are relatively infrequent. Third, lineages leave lakes more often than they enter lakes. For these reasons, riverine lineages have had greater opportunity to build diversity over long periods of time than lacustrine lineages.

## DISCUSSION

Why do freshwater habitats have higher species richness than expected given their total volume on Earth, especially compared to the ocean (Cohen 1970; Horn 1972)? Many past studies compared speciation and/or net diversification rates between freshwater and marine organisms in order to answer this question, but collectively have found mixed results (Table 1). In this study, I compared diversification rates in marine versus freshwater teleost fishes while separating freshwater species by riverine and lacustrine habitats. Contrary to expectations, marine and riverine fishes have similar net diversification rates on average. Lacustrine fishes have much faster diversification rates than all other aquatic habitats yet make up a small fraction of overall freshwater diversity. Therefore, I suggest that diversification rate differences are not sufficient to explain the freshwater paradox, and a more nuanced explanation is needed.

It is generally expected that freshwater fishes should have faster speciation and net diversification rates than marine fishes because freshwater habitats are more fragmented, so there are presumably more opportunities for speciation in freshwater than in the ocean (Grosberg et al. 2012; Tedesco et al. 2017a). This explanation implies that most freshwater diversity arose through allopatric speciation, an implication supported by the geographic distributions of closely related freshwater fish species (Lynch 1989; Hrbek et al. 2002; Near and Benard 2004; Dias et al. 2013; Albert et al. 2020; Sholihah et al. 2021). One potential problem with this explanation is that allopatric speciation is thought to proceed more slowly than sympatric speciation (McCune and Lovejoy 1998; Crow et al. 2010). Most freshwater diversity arose in rivers (Figs. 1, 3), which suggests that most freshwater species are indeed the product of allopatric speciation (assuming that this is the most common speciation mode in rivers). However, speciation and net diversification rates of riverine fishes were not much faster than those of marine fishes on average (Fig. 1). This finding seems to contradict the leading hypothesis for why freshwater fishes should have faster speciation rates than marine fishes. Still, this hypothesis is supported in part: allopatric speciation *sensu lato* appears to be important for explaining high freshwater species richness, but not necessarily fast rates of allopatric speciation. The fine-scale fragmentation of rivers might allow allopatric populations to complete the speciation process without being outcompeted by congeners or reabsorbed through hybridization (Germain et al. 2020), thus allowing the accumulation of many species over a relatively small spatial scale compared to other habitats. The density of speciation events in space can be decoupled from the frequency of events over time (Boucher et al. 2020).

How then can we explain the freshwater paradox, if not by faster diversification rates in freshwater versus the ocean? The results of biogeographic reconstructions (Fig. 3) suggest that explanations invoking geologic time could be more appropriate (Stephens and Wiens 2003; Wiens 2012). Diversity has been accumulating for comparably long periods of time in modern marine and riverine lineages of teleosts (over 100 million years; Fig. 3A). Given this amount of time, faster speciation is not necessary to explain high richness in freshwater. In fact, it seems that speciation is about equally as frequent in freshwater and the ocean (Fig. 1). This similarity in time-for-speciation, as well as the similarity in average net diversification rates, is consistent with the roughly similar species richness in riverine and marine habitats today (Fig. 1). Put another way, high freshwater diversity might be explained by the early colonization of freshwater in the history of teleosts, regular opportunities for speciation because of the continuous fragmentation of rivers, and protection from extinction allowing diversity to accumulate over long time scales. This time-based explanation for the freshwater paradox has also been suggested by past studies (Carrete Vega and Wiens 2012, Seehausen and Wagner 2014, Betancur-R et al. 2015; see also Miller and Wiens 2017, Miller and Román-Palacios 2021), but seems to have been given less attention from researchers than diversification rates (Table 1).

How do we reconcile the observation that speciation rates do not differ between riverine and marine habitats with evidence to the contrary from past studies? Manel et al. (2020) found that freshwater fishes have higher intraspecific genetic diversity than marine fishes. This presumably reflects the greater fragmentation of freshwater habitats, which could in principle lead to faster speciation rates. However, the authors found that genetic diversity was weakly related to local species richness (Correlation coefficient=0.20 in marine and 0.36 in freshwater, Manel et al. 2020). Population subdivision may not have a straightforward relationship with speciation rates since there may be distinct (and sometimes competing) factors driving population isolation, reproductive isolation, persistence and coexistence of incipient species (Harvey et al. 2019; Germain et al. 2020). For example, allopatric speciation creates daughter species with smaller population sizes than the parent species, which leads to greater risk of stochastic extinction (Moen and Morlon 2014). Further, species richness is itself only partially related to diversification rates and also depends on historical factors (Ricklefs 1987; Wiens 2012). Spatial patterns of species richness are more closely related to dispersal frequency and timing than to diversification rates in both the ocean (Miller et al. 2018) and freshwater (Miller and Román-Palacios 2021). Therefore, the links between population subdivision, speciation, and species richness can break down or be influenced by confounding factors.

Some macroevolutionary studies did find faster speciation and/or net diversification rates in freshwater (e.g., Bloom et al. 2013, Tedesco et al. 2017a, Miller et al. 2018; see Table 1). In light of the present study, it is possible that faster freshwater rates inferred by past studies were actually driven by lacustrine species, and not reflective of a general difference between marine and freshwater biomes. Indeed, Rabosky (2020) found that apparently faster average freshwater rates of speciation in actinopterygians were driven by select clades and were not broadly characteristic of marine-to-freshwater transitions. Lacustrine cichlids in particular have the fastest speciation rates among all fishes (Burress and Tan 2017; McGee et al. 2020). Future studies should aim to explain the variation in diversification rates seen within freshwater fishes, which are diverse in age and ecology.

The other side of the coin is that diversification rates in marine fishes are faster than expected given the apparent lack of barriers to dispersal in the ocean. Some marine lineages are known to have exceptionally fast speciation rates (Rüber and Zardoya 2005; Rabosky et al. 2018; Siqueira et al. 2020; see also Miller and Wiens 2017 for marine amniotes). Marine fishes are not a monolithic group, and a general expectation for low speciation rates in the ocean seems overly simplistic. The mechanisms for speciation in the marine environment appear to be as varied as those found in continental biomes, and speciation with gene flow may be common (Crow et al. 2010; Bowen et al. 2013). Furthermore, while there is a 7,500-fold difference in water volume between the ocean and freshwater (Horn 1972), most marine biodiversity is found along coastlines, not in the open ocean. In particular, coral reefs contain ∼40% of marine fishes within only 0.1% of the Earth’s surface area (Cowman et al. 2017). The similarity in species richness between marine and freshwater fishes (Fig. 1) does not seem as puzzling when considering the distribution of species within the ocean.

The most famous examples of lacustrine radiation are cichlids in the African rift lakes (Wagner et al. 2012; Seehausen 2015; Burress and Tan 2017; McGee et al. 2020). However, rapid diversification in lakes was not limited to cichlids (Fig. 2) nor African lakes (Fig. 3). Diversification rates were faster in lakes than in rivers within all six continental regions of the world (Fig. 3D). Lacustrine radiations with elevated diversification rates other than cichlids include: Lake Titicaca *Orestias* (mean DR=0.6 species/million years), *Coregonus* of Nearctic (DR=2.67) and Palearctic (2.09) postglacial lakes, Lake Baikal sculpins (0.71), Lake Tana *Labeobarbus* (4.08), and Sulawesi telmatherinid rainbowfishes (0.31). Additional examples are summarized in Table 4.

**Table 4.**
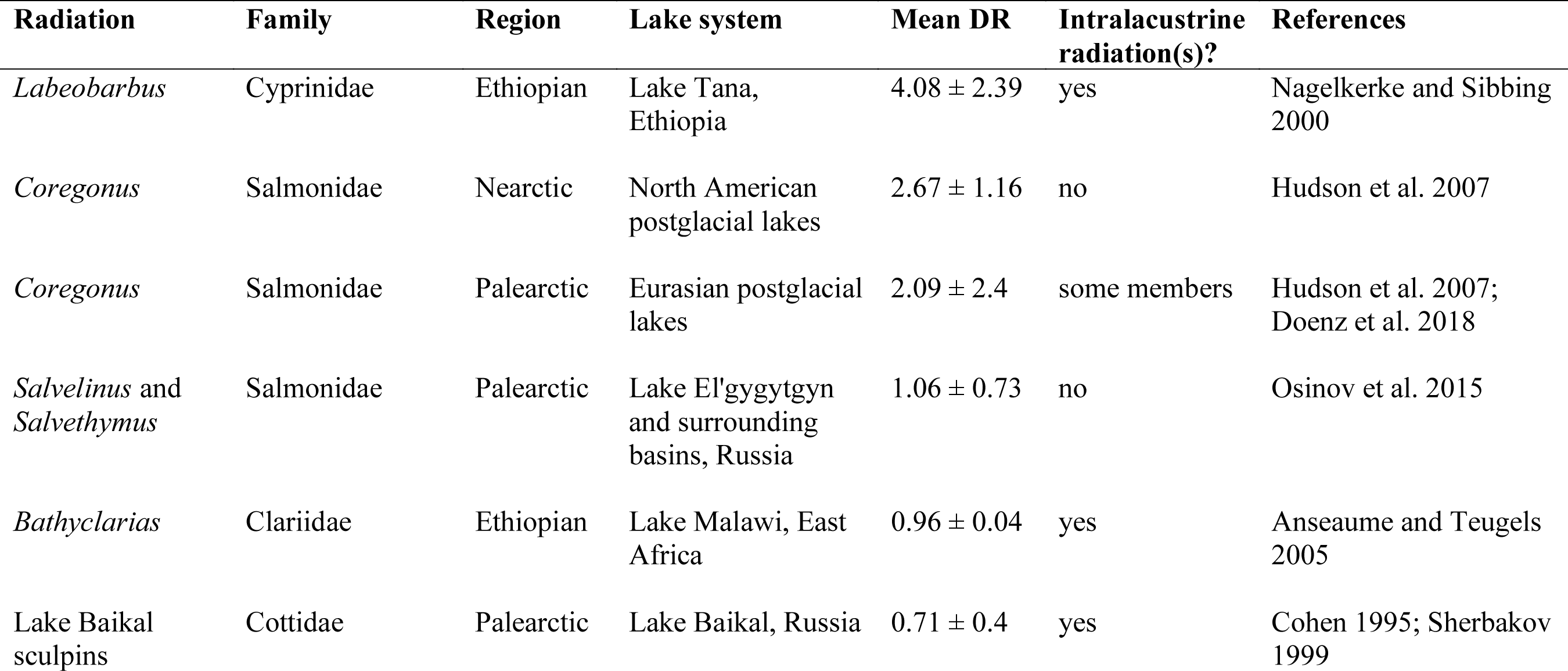
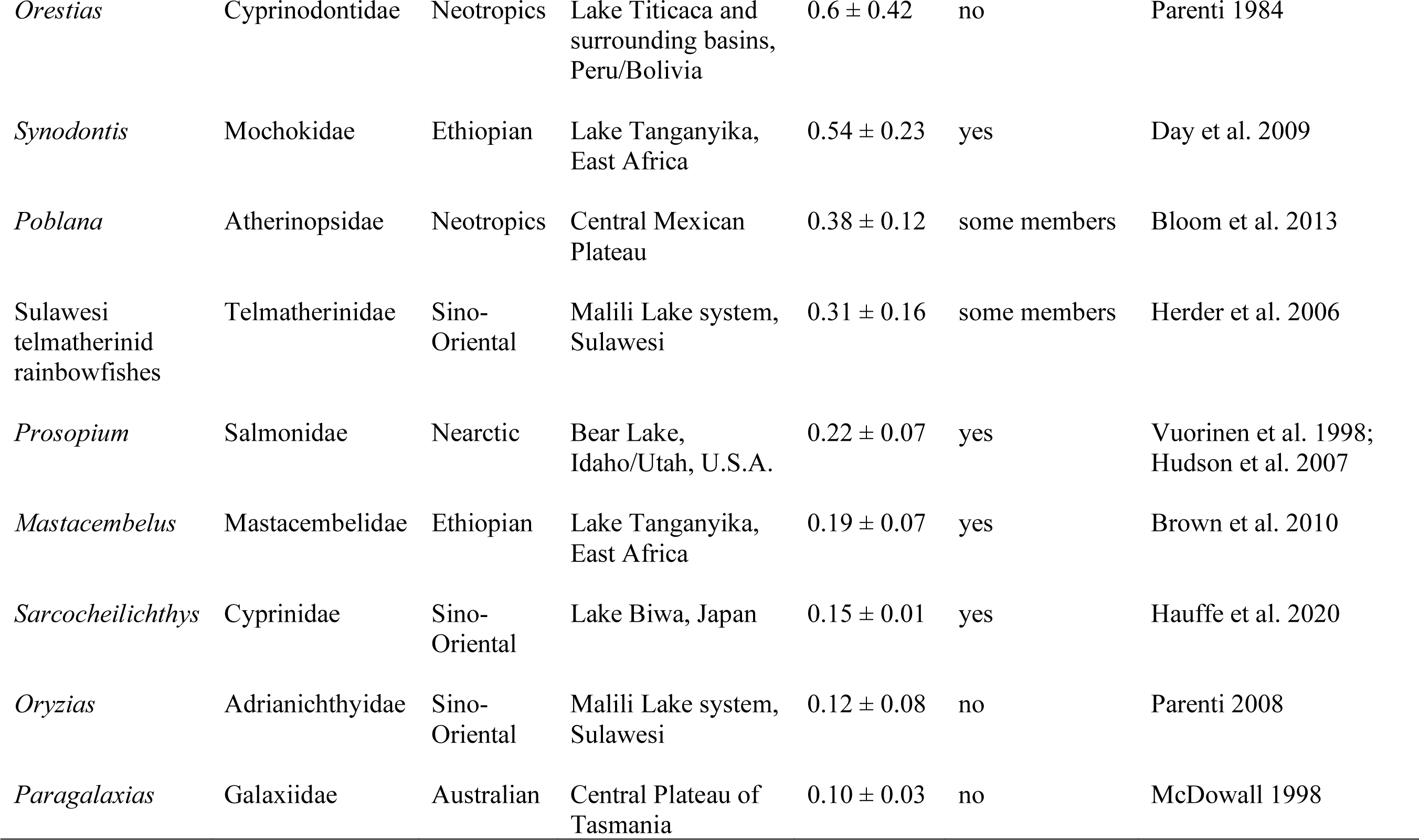
Examples of lacustrine radiations other than cichlids. Note that this list is not exhaustive. These radiations represent independent transitions to the lacustrine state based on biogeographic reconstructions (Fig. 3). Radiations are ranked by diversification rate in descending order. The mean DR represents the mean tip-associated diversification rate (species/million years) of members of the lacustrine lineage identified by BioGeoBEARS (Fig. 3D). Taxonomy follows Rabosky et al. (2018). Here I note whether monophyletic clade(s) are found within individual lakes (intralacustrine radiations) based on prior phylogenetic studies.

If diversification rates are so fast in lakes, then why do lakes contain a small proportion of freshwater diversity (Figs. 1, 3C)? Biogeographic models suggested that this contradictory pattern can be explained by transition rates among habitats (Fig. 3B). In the best-fitting model, direct transitions from the ocean into lakes were constrained to be rarer than marine-riverine transitions (Table 2). This is intuitive, as lakes are fed by rivers and do not usually interface directly with the ocean (but see Schluter and McPhail 1992 and Hughes et al. 2020 for examples of marine-lacustrine connections). The most common path to entering lakes is by first entering rivers, then occurring in both rivers and lakes, then finally becoming endemic to lakes (Fig. 3B). Since there are several evolutionary steps to becoming lacustrine, lacustrine radiations were usually nested within riverine lineages (Fig. 3A). Therefore, lacustrine radiations had less time-for-speciation than riverine counterparts (Fig. 3D; Stephens and Wiens 2003). In addition, transition rates out of lakes were higher than those into lakes (Fig. 3B). This suggests that being endemic to lakes is an evolutionarily unstable state.

These biogeographic results can be framed in light of the age of rivers versus lakes. On a local scale, conditions in rivers change frequently and seasonally, while conditions in lakes are more stable. However, lakes are generally younger than rivers over geologic timescales because lakes dry out and can disappear entirely (Lowe-McConnell 1969; McCune 1987; Joyce et al. 2005; Cristescu et al. 2010). Indeed, the fossil record documents past lacustrine radiations that went extinct and were replaced as lakes dried out and refilled (McCune 1987). Lacustrine fishes may avoid extinction by entering rivers (Joyce et al. 2005; Seehausen and Wagner 2014), explaining the high transition rates out of lakes. Another consideration is that rapid diversification rates are probably short lived, and the richness of lacustrine radiations may ultimately be constrained by the features of individual lakes (Wagner et al. 2012, 2014). It is noteworthy that many intralacustrine radiations are limited to populations within a single species, or pairs of species (McPhail and Schluter 1992, 1993; Hudson et al. 2007). In contrast, some large and ancient lakes host a greater diversity of species (Cohen 1995; Wagner et al. 2014) but these lake systems may be atypical.

It is somewhat unexpected that rivers have lower speciation rates than lakes, since the fragmentation of rivers promotes population subdivision and endemism (Dias et al. 2013). Note however that a speciation rate represents the number of species generated per unit of time, and not simply the number of species generated; therefore, the occurrence of many speciation events might not translate to a fast speciation rate if these events occurred over several million years. McCune and Lovejoy (1998) suggested that the reason for faster speciation rates in lakes was that sympatric speciation completes faster than allopatric speciation. My results across fishes are consistent with McCune and Lovejoy’s findings, assuming that sympatric speciation is more common in lakes than in rivers (Bolnick and Fitzpatrick 2007; Seehausen and Wagner 2014). Still, I did not directly identify geographic modes of speciation in this study. At the time of writing, phylogenetic sampling across fishes is still generally below that needed to accurately infer the speciation mode from present-day geographic ranges (Skeels and Cardillo 2019). In addition, allopatric speciation occurring over short distances within a lake can be difficult to distinguish from sympatric speciation without fine-scale spatial and ecological data (Rico and Turner 2002; Seehausen and van Alphen 1999). An unanswered question is whether allopatric speciation within and among lakes is also generally faster than allopatric speciation in rivers. Some lacustrine radiations for which allopatry appears to be the dominant mode also have rapid diversification rates (Table 4).

Besides speciation mode, there are ecological differences between rivers and lakes that could explain faster lacustrine diversification rates. Compared to rivers, lakes may allow for more ecological specialization because of their more stable conditions and greater structural diversity (Seehausen 2015). Stronger competition and predation pressure in rivers may also inhibit speciation (Briggs 1966; Lowe-McConnell 1969; Seehausen 2015), such that fishes experience ecological release when they colonize lakes (Yoder et al. 2010; Burress and Tan 2017). The observation that the present-day richness of many lakes appears to be below the lake’s carrying capacity (Barbour and Brown 1974; Doenz et al. 2018; Hauffe et al. 2020) supports the idea of high ecological opportunity in lakes.

In conclusion, I found that lacustrine fishes have faster speciation and net diversification rates than other aquatic habitats. Still, lakes represent a small fraction of freshwater diversity compared to rivers. Surprisingly, riverine and marine fishes had similar rates of diversification, despite past literature predicting that these habitats had opposite effects on diversification (Horn 1972; May 1994; Grosberg et al. 2012; Tedesco et al. 2017a). While allopatric speciation is indeed important for generating freshwater diversity, this may not be the mechanism driving faster speciation rates between freshwater and marine habitats observed in some past studies.

Furthermore, these results support an alternative explanation for the freshwater paradox. Speciation is roughly as frequent in freshwater and marine habitats on average, and teleosts have occupied these habitats for comparable amounts of time over their 200-million-year history.

Future research should aim to identify ecological and evolutionary mechanisms for how riverine habitats support long-term diversification and lineage persistence while lacustrine habitats promote rapid diversification.

## Acknowledgements

I thank John Wiens and Luke Tornabene for helpful discussion related to this manuscript. I was supported by an NSF Postdoctoral Fellowship (DBI-1906574).

## Author contributions

E.C.M. conceived the study, collected data, performed analyses, interpreted the results, and wrote the manuscript.

## Data Accessibility Statement

All data (including habitats for each species) and R code needed to replicate analyses are in the Dryad package associated with this manuscript. The temporary link to the repository during peer review is: XXXXXXX.

