## Supporting Information for "Comparing diversification rates in lakes, rivers, and the sea"

**In this file:**

Comparing rates among additional freshwater habitat types:
Extended Text 1: Details of additional freshwater habitat comparisons
Table S1: Co-distribution of additional habitats with rivers and lakes
Figure S1: Comparison of rates among additional freshwater habitat types

Full results of rate comparisons by habitat, original Rabosky et al. (2018) dates:
Table S2: Two habitats, all teleosts and cichlids removed
Table S3: Six habitats, all teleosts
Table S4: Four habitats, all teleosts
Table S5: Three habitats, all teleosts
Table S6: Six habitats, cichlids removed
Table S7: Four habitats, cichlids removed
Table S8: Three habitats, cichlids removed

Full results of rate comparisons by habitat using redated trees:
Table S9: Results of phylogenetic ANOVA using Alfaro et al. (2018) dates
Table S10: Results of phylogenetic ANOVA using Hughes et al. (2018) dates
Figure S2: Comparing rates between Rabosky et al. (2018) and redated trees

Biogeographic analyses:
Extended Text 2: Additional details of biogeographic models
Table S11: Full dispersal matrix used in biogeographic models
Extended Text 3: Assessing sensitivity to biased sampling

**Data availability:** All data (including habitat assignments for each species) and R code needed to replicate analyses are in the Dryad package associated with this manuscript (doi: XXXXXX).

**Extended Text 1: Details of additional freshwater habitat comparisons**

Some freshwater species were found in habitats that were not clearly lakes or rivers. One question is whether these habitats can be classified as either riverine or lacustrine in diversification rate comparisons, or if these habitats instead have distinct effects on diversification rates. While assigning habitat categories (based on FishBase or IUCN Red List descriptions of habitat), I noted if species occurred in the following five habitat types:

1. Wetlands including swamps, floodplains, marshes, bogs, fens, sloughs, and peats
2. Ponds
3. Pools, temporary pools, oxbow lakes, or billabongs
4. Caves
5. Springs

I found few species endemic to these habitats (18–22 species in each habitat; Table S1). Instead, many species found in these five habitats were also found in rivers, or both rivers and lakes (rarely lakes alone; Table S1). Due to the risk of phylogenetic pseudoreplication (Maddison and FitzJohn 2015), I did not compare diversification rates solely among the few species endemic to these habitats. Instead, I compared rates among habitats using all species found in each habitat type, including those also found in rivers and/or lakes. This was slightly different from my approach to comparing habitats in my main analyses (e.g. Figs. 1–2, Table 3). In those analyses, each species was assigned to a single category (rivers, lakes, or both rivers and lakes). To avoid comparing many categories with few species each, here I allowed each species to be classified under multiple habitat categories in the rate comparisons. This approach prevents statistical comparison of rates using phylogenetic ANOVA because species are counted multiple times. Instead, simply plotting the distribution of diversification rates found in each habitat should reveal if these habitats are likely to have distinct effects on diversification.

The distribution of diversification rates by habitat are shown in Figure S1 (DR statistic calculated without respect to traits from the phylogeny with all missing taxa imputed; Rabosky et al. 2018). There was much overlap in rates among the seven habitat categories (lakes, rivers, springs, caves, ponds, pools and wetlands; panel A). Rates were somewhat higher in lakes and springs, and somewhat lower in ponds and wetlands. Next, I separated species found solely in lakes from those co-distributed among lakes and the other habitats. Lake endemics had much faster rates than all other categories (panel B). This pattern remained after removing the family Cichlidae (panel C), though the mean rate for lake endemics was somewhat lower because lacustrine cichlids diversify at much faster rates than other fishes.

These results are summarized as follows. First, species endemic to habitats other than rivers or lakes are a very small portion of overall freshwater diversity. Second, species in these five additional habitats are usually also found in rivers, or rivers and lakes. Third, diversification rates of species found in these habitats are similar to those in the riverine or both rivers and lakes categories. The observations herein suggest that considering additional freshwater habitat categories will not change the key results reported in this study (principally that species endemic to lakes have faster diversification rates than other aquatic habitats).

For the purpose of the main analyses in this study, when species were found in these five additional habitat types, I used the species' distribution in rivers or lakes as the primary basis for its habitat categorization. I coded the few species endemic to other habitats as riverine, since these habitats are formed or maintained by fluvial action, they are most often co-distributed with rivers and their diversification rates are similar to rivers.

**Table S1.** Co-distribution of additional freshwater habitat types with rivers and lakes. Counts refer to species sampled in the phylogeny of Rabosky et al. (2018) that were found in these habitats See Extended Text 1 for more details.

| Habitat | Total number | Number endemic | Number shared with rivers | Number shared with rivers+lakes | Number shared with lakes |
|---|---|---|---|---|---|
| Wetlands | 362 | 19 | 197 | 119 | 9 |
| Ponds | 196 | 18 | 67 | 95 | 3 |
| Pools | 266 | 22 | 164 | 52 | 2 |
| Caves | 31 | 20 | 6 | 3 | 0 |
| Springs | 82 | 19 | 37 | 6 | 5 |

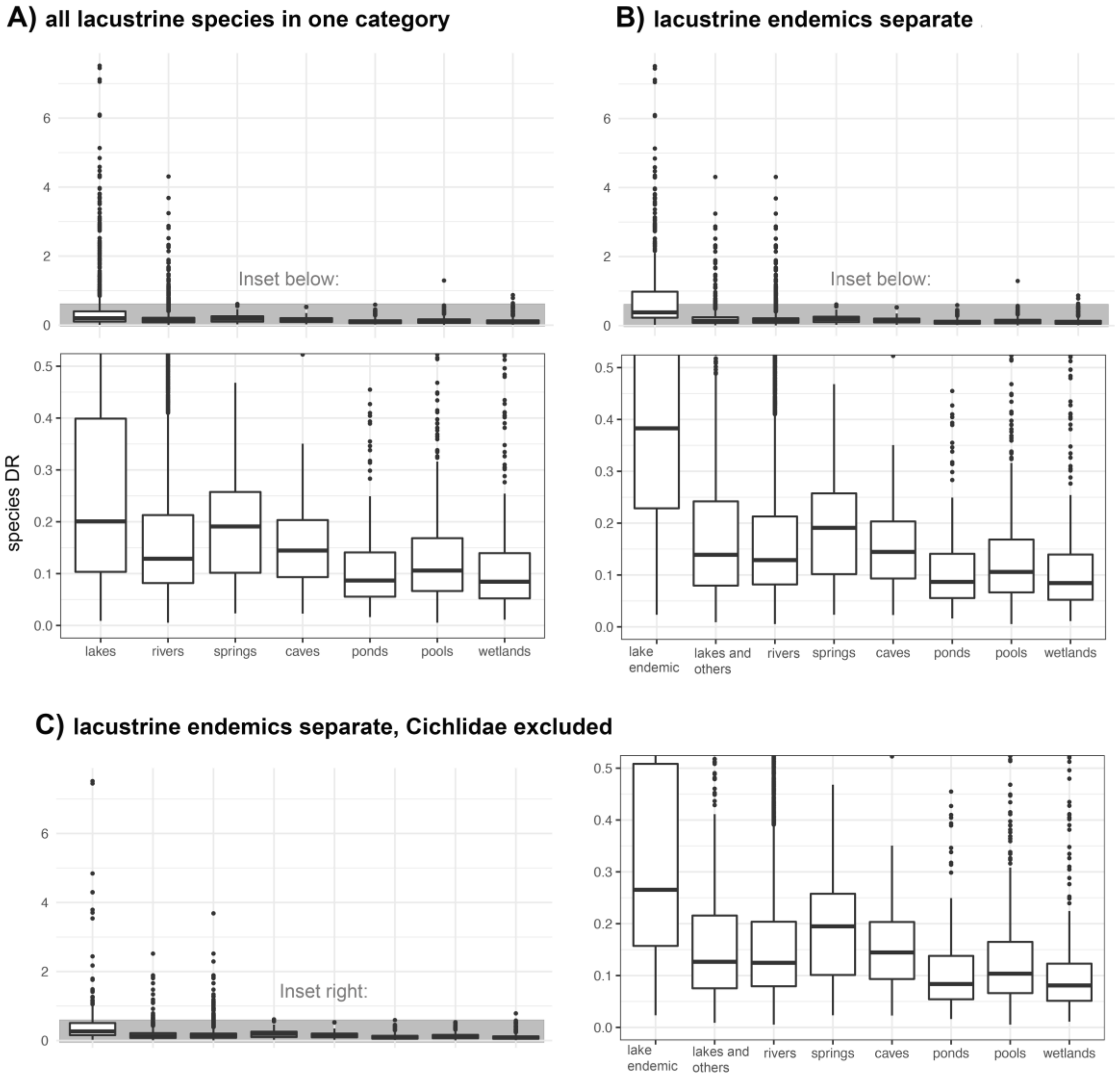


**Figure S1.** The distribution of diversification rates (DR statistic) found in freshwater habitat types (*n*=5,256 species). In panel A, categories represent all species in that habitat. Species can be found in multiple habitats. In panel B, species found only in lakes were separated from species found in lakes and other habitats. In panel C, lakes were treated as in panel B and cichlids were excluded. See Extended Text 1 and Table S1 for more details.

**Table S2.** Full results of diversification rate comparisons by habitat using phylogenetic ANOVA. Here I compared two habitat types (strictly marine versus strictly freshwater).

| | All teleosts (n=10,177) | | Cichlidae removed (n=9,457) | |
|---|---|---|---|---|
| Rate type | F | P-value | F | P-value |
| DR (all taxa added) | 203.69 | 0.653 | 68.82 | 0.789 |
| BAMM speciation, time-variable | 235.36 | 0.625 | 70.98 | 0.789 |
| BAMM speciation, time-constant | 285.95 | 0.537 | 109.27 | 0.747 |
| BAMM extinction, time-variable | 113.21 | 0.727 | 45.35 | 0.832 |
| BAMM extinction, time-constant | 207.37 | 0.635 | 102.59 | 0.748 |
| BAMM net div., time-variable | 230.59 | 0.624 | 59.59 | 0.811 |
| BAMM net div., time-constant | 250.59 | 0.597 | 67.23 | 0.793 |

**Table S3.** Full results of diversification rate comparisons by habitat using phylogenetic ANOVA including post hoc tests. Here I compared six habitat types and included all teleosts with phylogenetic data (n=11,055).

S3a. DR, all taxa added

F=441.86, P-value=0.093

Posthoc:

| | Lakes+rivers | Estuarine | Diadromous | Lacustrine | Marine | Riverine |
|---|---|---|---|---|---|---|
| Lakes+rivers | - | | | | | |
| Estuarine | 1.000 | - | | | | |
| Diadromous | 1.000 | 1.000 | - | | | |
| Lacustrine | **0.015** | **0.015** | **0.015** | - | | |
| Marine | 1.000 | 1.000 | 1.000 | **0.015** | - | |
| Riverine | 1.000 | 1.000 | 1.000 | **0.015** | 1.000 | - |

S3b. BAMM, speciation, time-variable model

F=551.71, P-value=0.062

Posthoc:

| | Lakes+rivers | Estuarine | Diadromous | Lacustrine | Marine | Riverine |
|---|---|---|---|---|---|---|
| Lakes+rivers | - | | | | | |
| Estuarine | 1.000 | - | | | | |
| Diadromous | 1.000 | 0.670 | - | | | |
| Lacustrine | **0.015** | **0.015** | **0.015** | - | | |
| Marine | 1.000 | 1.000 | 1.000 | **0.015** | - | |
| Riverine | 1.000 | 1.000 | 1.000 | **0.015** | 1.000 | - |

S3c. BAMM, speciation, time-constant model

F=564.22, P-value=0.059

Posthoc:

| | Lakes+rivers | Estuarine | Diadromous | Lacustrine | Marine | Riverine |
|---|---|---|---|---|---|---|
| Lakes+rivers | - | | | | | |
| Estuarine | 1.000 | - | | | | |
| Diadromous | 1.000 | 0.530 | - | | | |
| Lacustrine | **0.015** | **0.015** | **0.022** | - | | |
| Marine | 1.000 | 1.000 | 1.000 | **0.015** | - | |
| Riverine | 1.000 | 1.000 | 1.000 | **0.015** | 1.000 | - |

S3d. BAMM, extinction, time-variable model

F=231.26, P-value=0.298

Posthoc:

| | Lakes+rivers | Estuarine | Diadromous | Lacustrine | Marine | Riverine |
|---|---|---|---|---|---|---|
| Lakes+rivers | - | | | | | |
| Estuarine | 1.000 | - | | | | |
| Diadromous | 0.209 | **0.015** | - | | | |
| Lacustrine | 0.576 | 0.168 | 1.000 | - | | |
| Marine | 1.000 | 1.000 | **0.015** | 0.310 | - | |
| Riverine | 1.000 | 1.000 | 0.117 | 0.405 | 1.000 | - |

S3e. BAMM, extinction, time-constant model

F=272.27, P-value=0.27

Posthoc:

| | Lakes+rivers | Estuarine | Diadromous | Lacustrine | Marine | Riverine |
|---|---|---|---|---|---|---|
| Lakes+rivers | - | | | | | |
| Estuarine | 1.000 | - | | | | |
| Diadromous | 0.512 | **0.015** | - | | | |
| Lacustrine | 0.210 | **0.039** | 1.000 | - | | |
| Marine | 1.000 | 1.000 | **0.015** | 0.108 | - | |
| Riverine | 1.000 | 1.000 | 0.270 | 0.198 | 1.000 | - |

S3f. BAMM, net diversification, time-variable model

F=584.80, P-value=**0.047**

Posthoc:

| | Lakes+rivers | Estuarine | Diadromous | Lacustrine | Marine | Riverine |
|---|---|---|---|---|---|---|
| Lakes+rivers | - | | | | | |
| Estuarine | 1.000 | - | | | | |
| Diadromous | 1.000 | 1.000 | - | | | |
| Lacustrine | **0.015** | **0.015** | **0.015** | - | | |
| Marine | 1.000 | 1.000 | 1.000 | **0.015** | - | |
| Riverine | 1.000 | 1.000 | 1.000 | **0.015** | 1.000 | - |

S3g. BAMM, net diversification, time-constant model

F=598.17, P-value=**0.041**

Posthoc:

| | Lakes+rivers | Estuarine | Diadromous | Lacustrine | Marine | Riverine |
|---|---|---|---|---|---|---|
| Lakes+rivers | - | | | | | |
| Estuarine | 1.000 | - | | | | |
| Diadromous | 1.000 | 1.000 | - | | | |
| Lacustrine | **0.015** | **0.015** | **0.015** | - | | |
| Marine | 1.000 | 1.000 | 1.000 | **0.015** | - | |
| Riverine | 1.000 | 1.000 | 1.000 | **0.015** | 1.000 | - |

**Table S4.** Full results of diversification rate comparisons by habitat using phylogenetic ANOVA including post hoc tests. Here I compared four habitat types and included all teleosts with phylogenetic data (n=10,177).

S4a. DR, all taxa added

F=696.40, P-value=0.121

Posthoc:

| | Marine | Riverine | Lakes+rivers | Lacustrine |
|---|---|---|---|---|
| Marine | - | | | |
| Riverine | 1.000 | - | | |
| Lakes+rivers | 1.000 | 0.960 | - | |
| Lacustrine | **0.006** | **0.006** | **0.006** | - |

S4b. BAMM, speciation, time-variable model

F=858.03, P-value=0.071

Posthoc:

| | Marine | Riverine | Lakes+rivers | Lacustrine |
|---|---|---|---|---|
| Marine | - | | | |
| Riverine | 1.000 | - | | |
| Lakes+rivers | 1.000 | 0.621 | - | |
| Lacustrine | **0.006** | **0.006** | **0.006** | - |

S4c. BAMM, speciation, time-constant model

F=880.02, P-value=0.071

Posthoc:

| | Marine | Riverine | Lakes+rivers | Lacustrine |
|---|---|---|---|---|
| Marine | - | | | |
| Riverine | 1.000 | - | | |
| Lakes+rivers | 1.000 | 0.660 | - | |
| Lacustrine | **0.006** | **0.006** | **0.006** | - |

S4d. BAMM, extinction, time-variable model

F=307.70, P-value=0.344

Posthoc:

| | Marine | Riverine | Lakes+rivers | Lacustrine |
|---|---|---|---|---|
| Marine | - | | | |
| Riverine | 1.000 | - | | |
| Lakes+rivers | 1.000 | 0.462 | - | |
| Lacustrine | 0.078 | 0.110 | 0.112 | - |

S4e. BAMM, extinction, time-constant model

F=403.30, P-value=0.266

Posthoc:

| | Marine | Riverine | Lakes+rivers | Lacustrine |
|---|---|---|---|---|
| Marine | - | | | |
| Riverine | 1.000 | - | | |
| Lakes+rivers | 1.000 | 0.648 | - | |
| Lacustrine | **0.024** | **0.030** | **0.036** | - |

S4f. BAMM, net diversification, time-variable model

F=884.21, P-value=0.071

Posthoc:

| | Marine | Riverine | Lakes+rivers | Lacustrine |
|---|---|---|---|---|
| Marine | - | | | |
| Riverine | 1.000 | - | | |
| Lakes+rivers | 1.000 | 0.879 | - | |
| Lacustrine | **0.006** | **0.006** | **0.006** | - |

S4g. BAMM, net diversification, time-constant model

F=904.95, P-value=0.056

Posthoc:

| | Marine | Riverine | Lakes+rivers | Lacustrine |
|---|---|---|---|---|
| Marine | - | | | |
| Riverine | 1.000 | - | | |
| Lakes+rivers | 1.000 | 0.867 | - | |
| Lacustrine | **0.006** | **0.006** | **0.006** | - |

**Table S5.** Full results of diversification rate comparisons by habitat using phylogenetic ANOVA including post hoc tests. Here I compared three habitat types (freshwater only) and included all freshwater teleosts with phylogenetic data (n=5,256).

S5a. DR, all taxa added

F=588.28, P-value=**0.001**

Posthoc:

| | Riverine | Lakes+rivers | Lacustrine |
|---|---|---|---|
| Riverine | - | | |
| Lakes+rivers | 0.435 | - | |
| Lacustrine | **0.003** | **0.003** | - |

S5b. BAMM, speciation, time-variable model

F=722.03, P-value=**0.001**

Posthoc:

| | Riverine | Lakes+rivers | Lacustrine |
|---|---|---|---|
| Riverine | - | | |
| Lakes+rivers | 0.302 | - | |
| Lacustrine | **0.003** | **0.003** | - |

S5c. BAMM, speciation, time-constant model

F=726.23, P-value=**0.001**

Posthoc:

| | Riverine | Lakes+rivers | Lacustrine |
|---|---|---|---|
| Riverine | - | | |
| Lakes+rivers | 0.304 | - | |
| Lacustrine | **0.003** | **0.003** | - |

S5d. BAMM, extinction, time-variable model

F=260.83, P-value=0.061

Posthoc:

| | Riverine | Lakes+rivers | Lacustrine |
|---|---|---|---|

| | | | |
|---|---|---|---|
| Riverine | - | | |
| Lakes+rivers | 0.287 | - | |
| Lacustrine | 0.177 | 0.177 | - |

S5e. BAMM, extinction, time-constant model

F=726.23, P-value=**0.038**

Posthoc:

| | Riverine | Lakes+rivers | Lacustrine |
|---|---|---|---|
| Riverine | - | | |
| Lakes+rivers | 0.327 | - | |
| Lacustrine | 0.102 | 0.102 | - |

S5f. BAMM, net diversification, time-variable model

F=723.64, P-value=**0.001**

Posthoc:

| | Riverine | Lakes+rivers | Lacustrine |
|---|---|---|---|
| Riverine | - | | |
| Lakes+rivers | 0.451 | - | |
| Lacustrine | **0.003** | **0.003** | - |

S5g. BAMM, net diversification, time-constant model

F=746.08, P-value=**0.001**

Posthoc:

| | Riverine | Lakes+rivers | Lacustrine |
|---|---|---|---|
| Riverine | - | | |
| Lakes+rivers | 0.384 | - | |
| Lacustrine | **0.003** | **0.003** | - |

**Table S6.** Full results of diversification rate comparisons by habitat using phylogenetic ANOVA including post hoc tests. Here I compared six habitat types and discarded the family Cichlidae (n=10,331).

S6a. DR, all taxa added

F=129.04, P-value=0.464

Posthoc:

| | Lakes+rivers | Estuarine | Diadromous | Lacustrine | Marine | Riverine |
|---|---|---|---|---|---|---|
| Lakes+rivers | - | | | | | |
| Estuarine | 1.000 | - | | | | |
| Diadromous | 1.000 | 1.000 | - | | | |
| Lacustrine | **0.015** | **0.015** | **0.015** | - | | |
| Marine | 1.000 | 1.000 | 1.000 | **0.015** | - | |
| Riverine | 1.000 | 1.000 | 1.000 | **0.015** | 1.000 | - |

S6b. BAMM, speciation, time-variable model

F=140.72, P-value=0.428

Posthoc:

| | Lakes+rivers | Estuarine | Diadromous | Lacustrine | Marine | Riverine |
|---|---|---|---|---|---|---|
| Lakes+rivers | - | | | | | |
| Estuarine | 1.000 | - | | | | |
| Diadromous | 1.000 | 0.150 | - | | | |
| Lacustrine | **0.015** | **0.015** | **0.033** | - | | |
| Marine | 1.000 | 1.000 | 1.000 | **0.015** | - | |
| Riverine | 1.000 | 1.000 | 1.000 | **0.015** | 1.000 | - |

S6c. BAMM, speciation, time-constant model

F=150.18, P-value=0.418

Posthoc:

| | Lakes+rivers | Estuarine | Diadromous | Lacustrine | Marine | Riverine |
|---|---|---|---|---|---|---|
| Lakes+rivers | - | | | | | |
| Estuarine | 1.000 | - | | | | |
| Diadromous | 1.000 | 0.070 | - | | | |
| Lacustrine | **0.015** | **0.015** | **0.022** | - | | |
| Marine | 1.000 | 1.000 | 0.828 | **0.015** | - | |

| | | | | | | |
|---|---|---|---|---|---|---|
| Riverine | 1.000 | 1.000 | 1.000 | **0.015** | 1.000 | - |

S6d. BAMM, extinction, time-variable model

F=178.13, P-value=0.355

Posthoc:

| | Lakes+rivers | Estuarine | Diadromous | Lacustrine | Marine | Riverine |
|---|---|---|---|---|---|---|
| Lakes+rivers | - | | | | | |
| Estuarine | 1.000 | - | | | | |
| Diadromous | 0.144 | **0.015** | - | | | |
| Lacustrine | **0.015** | **0.015** | 1.000 | - | | |
| Marine | 1.000 | 1.000 | **0.015** | **0.020** | - | |
| Riverine | 1.000 | 1.000 | 0.072 | **0.015** | 1.000 | - |

S6e. BAMM, extinction, time-constant model

F=171.97, P-value=0.379

Posthoc:

| | Lakes+rivers | Estuarine | Diadromous | Lacustrine | Marine | Riverine |
|---|---|---|---|---|---|---|
| Lakes+rivers | - | | | | | |
| Estuarine | 1.000 | - | | | | |
| Diadromous | 0.384 | **0.015** | - | | | |
| Lacustrine | **0.015** | **0.015** | 1.000 | - | | |
| Marine | 1.000 | 1.000 | **0.015** | **0.015** | - | |
| Riverine | 1.000 | 1.000 | 0.324 | **0.015** | 1.000 | - |

S6f. BAMM, net diversification, time-variable model

F=93.15, P-value=0.555

Posthoc:

| | Lakes+rivers | Estuarine | Diadromous | Lacustrine | Marine | Riverine |
|---|---|---|---|---|---|---|
| Lakes+rivers | - | | | | | |
| Estuarine | 1.000 | - | | | | |
| Diadromous | 1.000 | 1.000 | - | | | |
| Lacustrine | **0.015** | **0.015** | **0.015** | - | | |
| Marine | 1.000 | 1.000 | 1.000 | **0.015** | - | |
| Riverine | 1.000 | 1.000 | 1.000 | **0.015** | 1.000 | - |

S6g. BAMM, net diversification, time-constant model

F=90.22, P-value=0.562

Posthoc:

| | Lakes+rivers | Estuarine | Diadromous | Lacustrine | Marine | Riverine |
|---|---|---|---|---|---|---|
| Lakes+rivers | - | | | | | |
| Estuarine | 1.000 | - | | | | |
| Diadromous | 1.000 | 1.000 | - | | | |
| Lacustrine | **0.015** | **0.015** | **0.015** | - | | |
| Marine | 1.000 | 1.000 | 1.000 | **0.015** | - | |
| Riverine | 1.000 | 1.000 | 1.000 | **0.015** | 1.000 | - |

**Table S7.** Full results of diversification rate comparisons by habitat using phylogenetic ANOVA including post hoc tests. Here I compared four habitat types and discarded the family Cichlidae (n=9,457).

S7a. DR, all taxa added

F=201.01, P-value=0.468

Posthoc:

| | Lakes+rivers | Lacustrine | Marine | Riverine |
|---|---|---|---|---|
| Lakes+rivers | - | | | |
| Lacustrine | **0.006** | - | | |
| Marine | 1.000 | **0.006** | - | |
| Riverine | 1.000 | **0.006** | 1.000 | - |

S7b. BAMM, speciation, time-variable model

F=197.73, P-value=0.448

Posthoc:

| | Lakes+rivers | Lacustrine | Marine | Riverine |
|---|---|---|---|---|
| Lakes+rivers | - | | | |
| Lacustrine | **0.006** | - | | |
| Marine | 1.000 | **0.006** | - | |
| Riverine | 1.000 | **0.006** | 1.000 | - |

S7c. BAMM, speciation, time-constant model

F=212.78, P-value=0.417

Posthoc:

| | Lakes+rivers | Lacustrine | Marine | Riverine |
|---|---|---|---|---|
| Lakes+rivers | - | | | |
| Lacustrine | **0.006** | - | | |
| Marine | 1.000 | **0.006** | - | |
| Riverine | 1.000 | **0.006** | 1.000 | - |

S7d. BAMM, extinction, time-variable model

F=188.51, P-value=0.487

Posthoc:

| | Lakes+rivers | Lacustrine | Marine | Riverine |
|---|---|---|---|---|
| Lakes+rivers | - | | | |
| Lacustrine | **0.006** | - | | |
| Marine | 1.000 | **0.006** | - | |
| Riverine | 0.789 | **0.006** | 1.000 | - |

S7e. BAMM, extinction, time-constant model

F=200.05, P-value=0.46

Posthoc:

| | Lakes+rivers | Lacustrine | Marine | Riverine |
|---|---|---|---|---|
| Lakes+rivers | - | | | |
| Lacustrine | **0.006** | - | | |
| Marine | 1.000 | **0.006** | - | |
| Riverine | 1.000 | **0.006** | 1.000 | - |

S7f. BAMM, net diversification, time-variable model

F=131.23, P-value=0.546

Posthoc:

| | Lakes+rivers | Lacustrine | Marine | Riverine |
|---|---|---|---|---|
| Lakes+rivers | - | | | |
| Lacustrine | **0.006** | - | | |
| Marine | 1.000 | **0.006** | - | |
| Riverine | 1.000 | **0.006** | 1.000 | - |

S7g. BAMM, net diversification, time-constant model

F=127.88, P-value=0.561

Posthoc:

| | Lakes+rivers | Lacustrine | Marine | Riverine |
|---|---|---|---|---|
| Lakes+rivers | - | | | |
| Lacustrine | **0.006** | - | | |
| Marine | 1.000 | **0.006** | - | |
| Riverine | 1.000 | **0.006** | 1.000 | - |

**Table S8.** Full results of diversification rate comparisons by habitat using phylogenetic ANOVA including post hoc tests. Here I compared three habitat types (freshwater only) and discarded the family Cichlidae (n=4,536).

S8a. DR, all taxa added

F=208.21, P-value=**0.001**

Posthoc:

| | Riverine | Lakes+rivers | Lacustrine |
|---|---|---|---|
| Riverine | - | | |
| Lakes+rivers | 0.792 | - | |
| Lacustrine | **0.003** | **0.003** | - |

S8b. BAMM, speciation, time-variable model

F=193.67, P-value=**0.001**

Posthoc:

| | Riverine | Lakes+rivers | Lacustrine |
|---|---|---|---|
| Riverine | - | | |
| Lakes+rivers | 0.575 | - | |
| Lacustrine | **0.003** | **0.003** | - |

S8c. BAMM, speciation, time-constant model

F=195.26, P-value=**0.001**

Posthoc:

| | Riverine | Lakes+rivers | Lacustrine |
|---|---|---|---|
| Riverine | - | | |
| Lakes+rivers | 0.641 | - | |
| Lacustrine | **0.003** | **0.003** | - |

S8d. BAMM, extinction, time-variable model

F=165.34, P-value=**0.001**

Posthoc:

| | Riverine | Lakes+rivers | Lacustrine |
|---|---|---|---|

| | | | |
|---|---|---|---|
| Riverine | - | | |
| Lakes+rivers | 0.379 | - | |
| Lacustrine | **0.003** | **0.003** | - |

S8e. BAMM, extinction, time-constant model

F=160.30, P-value=**0.001**

Posthoc:

| | Riverine | Lakes+rivers | Lacustrine |
|---|---|---|---|
| Riverine | - | | |
| Lakes+rivers | 0.576 | - | |
| Lacustrine | **0.003** | **0.003** | - |

S8f. BAMM, net diversification, time-variable model

F=122.68, P-value=**0.001**

Posthoc:

| | Riverine | Lakes+rivers | Lacustrine |
|---|---|---|---|
| Riverine | - | | |
| Lakes+rivers | 0.816 | - | |
| Lacustrine | **0.003** | **0.003** | - |

S8g. BAMM, net diversification, time-constant model

F=118.38, P-value=**0.001**

Posthoc:

| | Riverine | Lakes+rivers | Lacustrine |
|---|---|---|---|
| Riverine | - | | |
| Lakes+rivers | 0.806 | - | |
| Lacustrine | **0.003** | **0.003** | - |

**Table S9.** Full results of diversification rate comparisons by habitat using phylogenetic ANOVA including post hoc tests. Here I used a version of the Rabosky et al. (2018) phylogeny redated based on Alfaro et al. (2018). These analyses were performed using the clade Acanthomorpha alone, since this is the clade the two phylogenies have in common. All rate comparisons used the DR statistic calculated from the redated tree.

S9a. Two habitat comparison (n=6,423 species)

F=308.24, P-value=0.351

S9b. Two habitat comparison, Cichlidae excluded (n=6,423 species)

F=29.04, P-value=0.667

S9c. Six habitat comparison (n=7,060 species)

F=342.89, P-value=**0.031**

Posthoc:

| | Lakes+rivers | Estuarine | Diadromous | Lacustrine | Marine | Riverine |
|---|---|---|---|---|---|---|
| Lakes+rivers | - | | | | | |
| Estuarine | 1.000 | - | | | | |
| Diadromous | 1.000 | 1.000 | - | | | |
| Lacustrine | **0.015** | **0.026** | 0.066 | - | | |
| Marine | 1.000 | 1.000 | 1.000 | **0.036** | - | |
| Riverine | 0.510 | 1.000 | 1.000 | **0.015** | 1.000 | - |

S9d. Six habitat comparison, Cichlidae excluded (n=6,327 species)

F=48.31, P-value=0.271

Posthoc:

| | Lakes+rivers | Estuarine | Diadromous | Lacustrine | Marine | Riverine |
|---|---|---|---|---|---|---|
| Lakes+rivers | - | | | | | |
| Estuarine | 1.000 | - | | | | |
| Diadromous | 1.000 | 1.000 | - | | | |

| | | | | | | |
|---|---|---|---|---|---|---|
| Lacustrine | **0.015** | **0.015** | **0.022** | - | | |
| Marine | 1.000 | 1.000 | 1.000 | **0.015** | - | |
| Riverine | 0.510 | 1.000 | 1.000 | **0.015** | 1.000 | - |

S9e. Four habitat comparison (n=6,423 species)

F=509.27, P-value=0.052

Posthoc:

| | Marine | Riverine | Lakes+rivers | Lacustrine |
|---|---|---|---|---|
| Marine | - | | | |
| Riverine | 0.883 | - | | |
| Lakes+rivers | 0.880 | 0.195 | - | |
| Lacustrine | **0.032** | **0.006** | **0.006** | - |

S9f. Four habitat comparison, Cichlidae excluded (n=5,694 species)

F=60.23, P-value=0.297

Posthoc:

| | Marine | Riverine | Lakes+rivers | Lacustrine |
|---|---|---|---|---|
| Marine | - | | | |
| Riverine | 1.000 | - | | |
| Lakes+rivers | 1.000 | 1.000 | - | |
| Lacustrine | **0.012** | **0.006** | **0.006** | - |

S9g. Three habitat comparison (n=2,034 species)

F=223.45, P-value=**0.039**

Posthoc:

| | Riverine | Lakes+rivers | Lacustrine |
|---|---|---|---|
| Riverine | - | | |
| Lakes+rivers | 0.336 | - | |
| Lacustrine | 0.105 | 0.105 | - |

S9h. Three habitat comparison, Cichlidae excluded (n=2,034 species)

F=72.42, P-value=**0.001**

Posthoc:

| | Riverine | Lakes+rivers | Lacustrine |
|---|---|---|---|

| | | | |
|---|---|---|---|
| Riverine | - | | |
| Lakes+rivers | 0.844 | - | |
| Lacustrine | **0.003** | **0.003** | - |

**Table S10.** Full results of diversification rate comparisons by habitat using phylogenetic ANOVA including post hoc tests. Here I used a version of the Rabosky et al. (2018) phylogeny redated based on Hughes et al. (2018). These analyses were performed using the clade Teleostei. All rate comparisons used the DR statistic calculated from the redated tree.

S10a. Two habitat comparison (n=10,333 species)

F=186.19, P-value=0.731

S10b. Two habitat comparison, Cichlidae excluded (n=9,604 species)

F=82.56, P-value=0.833

S10c. Six habitat comparison (n=11,227 species)

F=445.18, P-value=0.219

Posthoc:

| | Lakes+rivers | Estuarine | Diadromous | Lacustrine | Marine | Riverine |
|---|---|---|---|---|---|---|
| Lakes+rivers | - | | | | | |
| Estuarine | 1.000 | - | | | | |
| Diadromous | 1.000 | 1.000 | - | | | |
| Lacustrine | **0.015** | **0.015** | **0.015** | - | | |
| Marine | 1.000 | 1.000 | 1.000 | **0.015** | - | |
| Riverine | 1.000 | 1.000 | 1.000 | **0.015** | 1.000 | - |

S10d. Six habitat comparison, Cichlidae excluded (n=10,494 species)

F=126.64, P-value=0.595

Posthoc:

| | Lakes+rivers | Estuarine | Diadromous | Lacustrine | Marine | Riverine |
|---|---|---|---|---|---|---|
| Lakes+rivers | - | | | | | |
| Estuarine | 1.000 | - | | | | |
| Diadromous | 1.000 | 1.000 | - | | | |
| Lacustrine | **0.015** | **0.015** | **0.015** | - | | |
| Marine | 1.000 | 1.000 | 1.000 | **0.015** | - | |

| | | | | | | |
|---|---|---|---|---|---|---|
| Riverine | 1.000 | 1.000 | 1.000 | **0.015** | 1.000 | - |

S10e. Four habitat comparison (n=10,333 species)

F=691.60, P-value=0.259

Posthoc:

| | Marine | Riverine | Lakes+rivers | Lacustrine |
|---|---|---|---|---|
| Marine | - | | | |
| Riverine | 1.000 | - | | |
| Lakes+rivers | 1.000 | 0.876 | - | |
| Lacustrine | **0.006** | **0.010** | **0.010** | - |

S10f. Four habitat comparison, Cichlidae excluded (n=9,604 species)

F=199.39, P-value=0.560

Posthoc:

| | Marine | Riverine | Lakes+rivers | Lacustrine |
|---|---|---|---|---|
| Marine | - | | | |
| Riverine | 1.000 | - | | |
| Lakes+rivers | 1.000 | 1.000 | - | |
| Lacustrine | **0.006** | **0.006** | **0.006** | - |

S10g. Three habitat comparison (n=5,349 species)

F=537.31, P-value=**0.001**

Posthoc:

| | Riverine | Lakes+rivers | Lacustrine |
|---|---|---|---|
| Riverine | - | | |
| Lakes+rivers | 0.404 | - | |
| Lacustrine | **0.003** | **0.004** | - |

S10h. Three habitat comparison, Cichlidae excluded (n=4,620 species)

F=188.64, P-value=**0.001**

Posthoc:

| | Riverine | Lakes+rivers | Lacustrine |
|---|---|---|---|
| Riverine | - | | |
| Lakes+rivers | 0.552 | - | |

| | | | |
|---|---|---|---|
| Lacustrine | **0.003** | **0.003** | **-** |

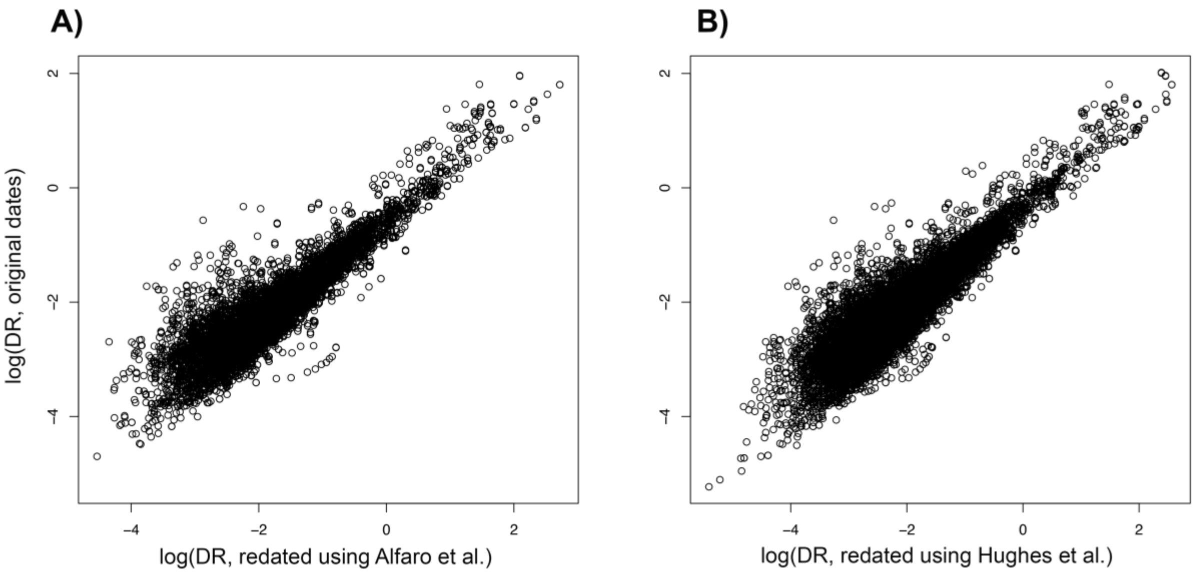


**Figure S2.** Comparison of diversification rates (DR statistic) between the original divergence times of Rabosky et al. (2018) and divergence times resulting from congruification using Alfaro et al. (2018; panel A) or Hughes et al. (2018; panel B) as a reference phylogeny (see Eastman et al. 2013 for congruification method). Note that rates using the Alfaro et al. dates represent the clade Acanthomorpha alone; rates using the Hughes et al. dates represent the clade Teleostei.

**Extended Text 2: Additional details of biogeographic models**

Here I describe in greater detail how I set up biogeographic model fitting. I fit 36 alternative biogeographic models using BioGeoBEARS version 1.1.2 (Matzke 2014; see Table 2). The details in this section are shared by all models.

I modeled dispersal among 13 habitat-region combinations. I included six biogeographic regions following Leroy et al. (2019), who used clustering algorithms to identify a bioregionalization scheme based on shared freshwater fish diversity. These were: (1) Nearctic, including North America including northern Mexico, (2) Neotropical, or southern Mexico, Central and South America; (3) Palearctic, or Europe, most of the Arabian Peninsula, western Asia, and Siberia; (4) Ethiopian, or the African continent and parts of the Arabian Peninsula; (5) Sino-Oriental, or Southern and Eastern Asia spanning India, China, Korea, Japan, the Sunda Shelf and associated islands; and (6) Australia including Papua New Guinea. I modified this regionalization slightly by including the Caribbean islands in the Neotropical region, including Madagascar in the Ethiopian region, and including islands of Oceania in the Australian region. Of 5,349 freshwater species in the Rabosky et al. (2018) phylogeny, 4,435 species (82.9%) were assigned to regions using the occurrence record database of Tedesco et al. (2017) updated by Leroy et al. (2019). The remaining 914 species not found in that database were assigned to regions based on descriptions of species' geographic range in FishBase or the IUCN Red List.

Species in these six regions were further assigned to riverine, lacustrine, or both habitat categories for a total of 12 freshwater region-habitat combinations. Species found in marine and estuarine habitats were assigned to a $13^{th}$ "marine" region. Diadromous species or those with unclear habitat affinities were excluded from biogeographic models. The computational intensiveness of biogeographic models scales exponentially with the number of possible ranges that can be reconstructed at a given node (Matzke 2014). To keep the model fitting feasible at this very large phylogenetic scale, I limited the maximum range size to two. Such combinations could include a species found in a single region in both lakes and rivers, two regions in rivers only, but not two regions in both lakes and rivers. Most species were endemic to a single region or habitat type: I excluded only 80 species that were found in two regions and both habitat types from biogeographic analyses. In total, 10,924 species were included in these analyses. I restricted the root of the teleost phylogeny to the marine state in accordance with the fossil record of fishes (Betancur-R et al. 2015).

All models were time-stratified in accordance with changing connectivity of regions through the ~200-million-year history of teleosts. To maintain consistency with prior literature, I followed DEC analyses from Toussaint et al. (2017) for freshwater beetles and Miller and Román-Palacios (2021) for freshwater fishes. The following rules were applied to all five time bins. Dispersal between adjacent regions was not constrained (i.e. the probability of dispersal between adjacent regions was multiplied by 1). The probability of dispersal between regions separated by a small marine barrier was multiplied by 0.75. The probability of dispersal between regions separated by another landmass was multiplied by 0.50. Finally, dispersal between regions separated by a large marine barrier was multiplied by 0.25. See full details of time bins and relevant geologic events in Table S11.

I also applied constraints on habitat transitions in all models. The transition probability between marine and freshwater habitats was multiplied by 0.25 at all times. Instantaneous region-habitat transitions were disallowed: for example, lineages could not directly disperse from Neotropical rivers to Nearctic lakes. Transitions between freshwater habitats within the same region were unconstrained (e.g. there were no constraints on the probability of transitioning from Neotropical rivers to Neotropical lakes).

All other details are given in the main text, Table 2, and Table S11.

**Table S11.** Baseline dispersal matrix between regions and habitats used in time-stratified biogeographic model fitting (Matzke, 2014). We constrained dispersal probabilities with time in concordance with tectonic activity and changing connectivity among regions. That is, dispersal between adjacent regions was not constrained; dispersal probability among regions separated by a small marine barrier was set to 0.75; dispersal probability among regions separated by another landmass was set to 0.50; and dispersal probability among regions separated by a large marine barrier was set to 0.25. The transition probability between marine and freshwater habitats was set to 0.25 at all times. We made further modifications to these baseline rules to test hypotheses about the role of lakes in dispersal (Table 2). In models 2, 5 and 6 transitions between marine and lake habitats (**orange values**) were lowered to 0.05. In models 4 and 6 dispersals between regions via lakes were lowered to 0.05 if they take place across long overseas or landmass barriers (**purple values**). In models 3 and 5 all dispersals between regions via lakes were lowered to 0.05 (**purple** and **green** values). Here region names and boundaries follow Leroy et al. (2019): Neo=Neotropical, Pal=Palearctic, Near=Nearctic, Eth=Ethiopian, Aust=Australia, SinO=Sino-Oriental. Habitat abbreviations are: Mar=marine, R=riverine, and L=lacustrine.

**S11a. 0–20 mya.** Nearctic and Neotropic regions connected by Isthmus of Panama; Tethys Ocean closed connecting Africa and Europe; Nearctic and Europe intermittently connected

| | Mar | Neo-R | Pal-R | Near-R | Eth-R | Aust-R | SinO-R | Neo-L | Pal-L | Near-L | Eth-L | Aust-L | SinO-L |
|---|---|---|---|---|---|---|---|---|---|---|---|---|---|
| Mar | - | 0.25 | 0.25 | 0.25 | 0.25 | 0.25 | 0.25 | **0.25** | **0.25** | **0.25** | **0.25** | **0.25** | **0.25** |
| Neo-R | 0.25 | - | 0.25 | 1 | 0.25 | 0.25 | 0.25 | 1 | 0 | 0 | 0 | 0 | 0 |
| Pal-R | 0.25 | 0.25 | - | 0.75 | 1 | 0.5 | 1 | 0 | 1 | 0 | 0 | 0 | 0 |
| Near-R | 0.25 | 1 | 0.75 | - | 0.25 | 0.25 | 0.25 | 0 | 0 | 1 | 0 | 0 | 0 |
| Eth-R | 0.25 | 0.25 | 1 | 0.25 | - | 0.25 | 0.5 | 0 | 0 | 0 | 1 | 0 | 0 |
| Aust-R | 0.25 | 0.25 | 0.5 | 0.25 | 0.25 | - | 1 | 0 | 0 | 0 | 0 | 1 | 0 |
| SinO-R | 0.25 | 0.25 | 1 | 0.25 | 0.5 | 1 | - | 0 | 0 | 0 | 0 | 0 | 1 |

| | | | | | | | | | | | | |
|---|---|---|---|---|---|---|---|---|---|---|---|---|
| Neo-L | **0.25** | 1 | 0 | 0 | 0 | 0 | 0 | - | **0.25** | **1** | **0.25** | **0.25** | **0.25** |
| Pal-L | **0.25** | 0 | 1 | 0 | 0 | 0 | 0 | **0.25** | - | **0.75** | **1** | **0.5** | **1** |
| Near-L | **0.25** | 0 | 0 | 1 | 0 | 0 | 0 | **1** | **0.75** | - | **0.25** | **0.25** | **0.25** |
| Eth-L | **0.25** | 0 | 0 | 0 | 1 | 0 | 0 | **0.25** | **1** | **0.25** | - | **0.25** | **0.5** |
| Aust-L | **0.25** | 0 | 0 | 0 | 0 | 1 | 0 | **0.25** | **0.5** | **0.25** | **0.25** | - | **1** |
| SinO-L | **0.25** | 0 | 0 | 0 | 0 | 0 | 1 | **0.25** | **1** | **0.25** | **0.5** | **1** | - |

**S11b. 20–40 mya.** India closer to Africa via Arabian Peninsula; Australia approaching Indo-Malay; Isthmus of Panama not connected; Africa and Europe separated by Tethys Ocean; Europe and Nearctic connected through Beringia

| | Mar | Neo-R | Pal-R | Near-R | Eth-R | Aust-R | SinO-R | Neo-L | Pal-L | Near-L | Eth-L | Aust-L | SinO-L |
|---|---|---|---|---|---|---|---|---|---|---|---|---|---|
| Mar | - | 0.25 | 0.25 | 0.25 | 0.25 | 0.25 | 0.25 | **0.25** | **0.25** | **0.25** | **0.25** | **0.25** | **0.25** |
| Neo-R | 0.25 | - | 0.25 | 0.75 | 0.25 | 0.25 | 0.25 | 1 | 0 | 0 | 0 | 0 | 0 |
| Pal-R | 0.25 | 0.25 | - | 1 | 0.75 | 0.25 | 1 | 0 | 1 | 0 | 0 | 0 | 0 |
| Near-R | 0.25 | 0.75 | 1 | - | 0.25 | 0.25 | 0.25 | 0 | 0 | 1 | 0 | 0 | 0 |
| Eth-R | 0.25 | 0.25 | 0.75 | 0.25 | - | 0.25 | 0.75 | 0 | 0 | 0 | 1 | 0 | 0 |
| Aust-R | 0.25 | 0.25 | 0.25 | 0.25 | 0.25 | - | 0.25 | 0 | 0 | 0 | 0 | 1 | 0 |
| SinO-R | 0.25 | 0.25 | 1 | 0.25 | 0.75 | 0.25 | - | 0 | 0 | 0 | 0 | 0 | 1 |
| Neo-L | **0.25** | 1 | 0 | 0 | 0 | 0 | 0 | - | **0.25** | **0.75** | **0.25** | **0.25** | **0.25** |
| Pal-L | **0.25** | 0 | 1 | 0 | 0 | 0 | 0 | **0.25** | - | **1** | **0.75** | **0.25** | **1** |
| Near-L | **0.25** | 0 | 0 | 1 | 0 | 0 | 0 | **0.75** | **1** | - | **0.25** | **0.25** | **0.25** |
| Eth-L | **0.25** | 0 | 0 | 0 | 1 | 0 | 0 | **0.25** | **0.75** | **0.25** | - | **0.25** | **0.75** |
| Aust-L | **0.25** | 0 | 0 | 0 | 0 | 1 | 0 | **0.25** | **0.25** | **0.25** | **0.25** | - | **0.25** |
| SinO-L | **0.25** | 0 | 0 | 0 | 0 | 0 | 1 | **0.25** | **1** | **0.25** | **0.75** | **0.25** | - |

**S11c. 40-80 mya.** South America and Africa are separated; Australia still connected to Antarctica

| | Mar | Neo-R | Pal-R | Near-R | Eth-R | Aust-R | SinO-R | Neo-L | Pal-L | Near-L | Eth-L | Aust-L | SinO-L |
|---|---|---|---|---|---|---|---|---|---|---|---|---|---|
| Mar | - | 0.25 | 0.25 | 0.25 | 0.25 | 0.25 | 0.25 | **0.25** | **0.25** | **0.25** | **0.25** | **0.25** | **0.25** |
| Neo-R | 0.25 | - | 0.25 | 0.75 | 0.75 | 0.5 | 0.25 | 1 | 0 | 0 | 0 | 0 | 0 |
| Pal-R | 0.25 | 0.25 | - | 1 | 0.75 | 0.25 | 1 | 0 | 1 | 0 | 0 | 0 | 0 |
| Near-R | 0.25 | 0.75 | 1 | - | 0.25 | 0.25 | 0.25 | 0 | 0 | 1 | 0 | 0 | 0 |

| | | | | | | | | | | | | |
|---|---|---|---|---|---|---|---|---|---|---|---|---|
| Eth-R | 0.25 | 0.75 | 0.75 | 0.25 | - | 0.25 | 0.75 | 0 | 0 | 0 | 1 | 0 | 0 |
| Aust-R | 0.25 | 0.5 | 0.25 | 0.25 | 0.25 | - | 0.25 | 0 | 0 | 0 | 0 | 1 | 0 |
| SinO-R | 0.25 | 0.25 | 1 | 0.25 | 0.75 | 0.25 | - | 0 | 0 | 0 | 0 | 0 | 1 |
| Neo-L | **0.25** | 1 | 0 | 0 | 0 | 0 | 0 | - | **0.25** | **0.75** | **0.75** | **0.5** | **0.25** |
| Pal-L | **0.25** | 0 | 1 | 0 | 0 | 0 | 0 | **0.25** | - | **1** | **0.75** | **0.25** | **1** |
| Near-L | **0.25** | 0 | 0 | 1 | 0 | 0 | 0 | **0.75** | **1** | - | **0.25** | **0.25** | **0.25** |
| Eth-L | **0.25** | 0 | 0 | 0 | 1 | 0 | 0 | **0.75** | **0.75** | **0.25** | - | **0.25** | **0.75** |
| Aust-L | **0.25** | 0 | 0 | 0 | 0 | 1 | 0 | **0.5** | **0.25** | **0.25** | **0.25** | - | **0.25** |
| SinO-L | **0.25** | 0 | 0 | 0 | 0 | 0 | 1 | **0.25** | **1** | **0.25** | **0.75** | **0.25** | - |

**S11d. 80–150 mya.** South America and Africa connected; India and Palearctic connected to Africa by land

| | Mar | Neo-R | Pal-R | Near-R | Eth-R | Aust-R | SinO-R | Neo-L | Pal-L | Near-L | Eth-L | Aust-L | SinO-L |
|---|---|---|---|---|---|---|---|---|---|---|---|---|---|
| Mar | - | 0.25 | 0.25 | 0.25 | 0.25 | 0.25 | 0.25 | **0.25** | **0.25** | **0.25** | **0.25** | **0.25** | **0.25** |
| Neo-R | 0.25 | - | 0.5 | 0.75 | 1 | 0.5 | 0.5 | 1 | 0 | 0 | 0 | 0 | 0 |
| Pal-R | 0.25 | 0.5 | - | 1 | 1 | 0.25 | 1 | 0 | 1 | 0 | 0 | 0 | 0 |
| Near-R | 0.25 | 0.75 | 1 | - | 0.25 | 0.25 | 0.25 | 0 | 0 | 1 | 0 | 0 | 0 |
| Eth-R | 0.25 | 1 | 1 | 0.25 | - | 0.25 | 0.75 | 0 | 0 | 0 | 1 | 0 | 0 |
| Aust-R | 0.25 | 0.5 | 0.25 | 0.25 | 0.25 | - | 0.75 | 0 | 0 | 0 | 0 | 1 | 0 |
| SinO-R | 0.25 | 0.5 | 1 | 0.25 | 0.75 | 0.75 | - | 0 | 0 | 0 | 0 | 0 | 1 |
| Neo-L | **0.25** | 1 | 0 | 0 | 0 | 0 | 0 | - | **0.5** | **0.75** | **1** | **0.5** | **0.5** |
| Pal-L | **0.25** | 0 | 1 | 0 | 0 | 0 | 0 | **0.5** | - | **1** | **1** | **0.25** | **1** |
| Near-L | **0.25** | 0 | 0 | 1 | 0 | 0 | 0 | **0.75** | **1** | - | **0.25** | **0.25** | **0.25** |
| Eth-L | **0.25** | 0 | 0 | 0 | 1 | 0 | 0 | **1** | **1** | **0.25** | - | **0.25** | **0.75** |
| Aust-L | **0.25** | 0 | 0 | 0 | 0 | 1 | 0 | **0.5** | **0.25** | **0.25** | **0.25** | - | **0.75** |
| SinO-L | **0.25** | 0 | 0 | 0 | 0 | 0 | 1 | **0.5** | **1** | **0.25** | **0.75** | **0.75** | - |

**S11e. 150–193 mya (root of teleost phylogeny).** Pangaea was intact

| | Mar | Neo-R | Pal-R | Near-R | Eth-R | Aust-R | SinO-R | Neo-L | Pal-L | Near-L | Eth-L | Aust-L | SinO-L |
|---|---|---|---|---|---|---|---|---|---|---|---|---|---|
| Mar | - | 0.25 | 0.25 | 0.25 | 0.25 | 0.25 | 0.25 | **0.25** | **0.25** | **0.25** | **0.25** | **0.25** | **0.25** |
| Neo-R | 0.25 | - | 0.5 | 1 | 1 | 0.5 | 0.5 | 1 | 0 | 0 | 0 | 0 | 0 |
| Pal-R | 0.25 | 0.5 | - | 1 | 0.5 | 0.25 | 1 | 0 | 1 | 0 | 0 | 0 | 0 |

| | | | | | | | | | | | | | |
|---|---|---|---|---|---|---|---|---|---|---|---|---|---|
| Near-R | 0.25 | 1 | 1 | - | 1 | 0.25 | 0.25 | 0 | 0 | 1 | 0 | 0 | 0 |
| Eth-R | 0.25 | 1 | 0.5 | 1 | - | 0.5 | 1 | 0 | 0 | 0 | 1 | 0 | 0 |
| Aust-R | 0.25 | 0.5 | 0.25 | 0.25 | 0.5 | - | 1 | 0 | 0 | 0 | 0 | 1 | 0 |
| SinO-R | 0.25 | 0.5 | 1 | 0.25 | 1 | 1 | - | 0 | 0 | 0 | 0 | 0 | 1 |
| Neo-L | **0.25** | 1 | 0 | 0 | 0 | 0 | 0 | **-** | **0.5** | **1** | **1** | **0.5** | **0.5** |
| Pal-L | **0.25** | 0 | 1 | 0 | 0 | 0 | 0 | **0.5** | **-** | **1** | **0.5** | **0.25** | **1** |
| Near-L | **0.25** | 0 | 0 | 1 | 0 | 0 | 0 | **1** | **1** | **-** | **1** | **0.25** | **0.25** |
| Eth-L | **0.25** | 0 | 0 | 0 | 1 | 0 | 0 | **1** | **0.5** | **1** | **-** | **0.5** | **1** |
| Aust-L | **0.25** | 0 | 0 | 0 | 0 | 1 | 0 | **0.5** | **0.25** | **0.25** | **0.5** | **-** | **1** |
| SinO-L | **0.25** | 0 | 0 | 0 | 0 | 0 | 1 | **0.5** | **1** | **0.25** | **1** | **1** | **-** |

**Extended Text 3: Assessing sensitivity to biased sampling**

The richness of independent transition events shown in Figure 3D was inferred by counting tips in the phylogeny. This was necessary because I could not assign species not sampled in the phylogeny to individual habitat transition events in the absence of a biogeographic ancestral reconstruction. I found that richness-per-transition was higher in rivers than lakes for most biogeographic regions.

The purpose of this section is to assess how this result might be influenced by sampling biases among habitats (i.e., if rivers were better sampled than lakes in the phylogeny). The proportion of described teleost species sampled in the phylogeny is known (Rabosky et al. 2018). While I do not know the distribution of unsampled species found in each habitat, I can identify taxonomic groups that are mostly marine, mostly riverine, and mostly lacustrine based on representatives in my dataset. Then, I can assess if these groups differ in their species sampling. This approach assumes that if species sampled for a clade are mostly found in one habitat, then the unsampled species in that genus are likely to share that habitat.

I assigned genera to one of three habitat groups (mostly or entirely marine, mostly or entirely riverine, and mostly or entirely lacustrine). I used a cut-off of 75% of representative species found in each habitat to assign genera to these categories. I did not include monotypic genera (or genera with only 1 member sampled in the tree) because these may not reveal much about sampling completeness at the level of species. I also did not assess sampling bias for the "both riverine and lacustrine" category because my biogeographic analyses strongly suggest that this state is transitional and is not likely to be conserved across a single genus (Figure 3).

I identified 733 marine genera, 407 riverine genera, and 52 lacustrine genera. The mean proportion of species sampled among marine genera was 59.9% (standard deviation ±30.3%).

The mean proportion of species sampled among riverine genera was 57.8% (standard deviation ±31.7%). The mean proportion of species sampled among lacustrine genera was 70.2% (standard deviation ±33.5%). These proportions suggest that phylogenetic sampling is highest for lacustrine genera, and similar for marine and riverine genera. This means that the low richness-per-transition to lacustrine habitats is not attributable to lower sampling of lacustrine species in this phylogeny relative to rivers (Fig. 3D).

Note that I also found that lacustrine diversification rates were higher than marine or riverine rates (Figs. 1–3). Tip-associated diversification rates can also be biased by sampling, because greater sampling decreases the distance between nodes which may imitate faster diversification. However, my results using the DR statistic calculated from the tree with all unsampled species grafted to the phylogeny (see details in Rabosky et al. 2018, Chang et al. 2019) were very similar to rates calculated from BAMM using the tree including species with genetic data only (Table 3, Tables S2–S8). Note also that BAMM corrects for incomplete sampling on a per-taxon basis. Therefore, it is unlikely that faster diversification rates in lacustrine habitats is an artefact of greater sampling of lacustrine species.
